# TWO-SIGMA-G: A New Competitive Gene Set Testing Framework for scRNA-seq Data Accounting for Inter-Gene and Cell-Cell Correlation

**DOI:** 10.1101/2021.01.24.427979

**Authors:** Eric Van Buren, Ming Hu, Liang Cheng, John Wrobel, Kirk Wilhelmsen, Lishan Su, Yun Li, Di Wu

## Abstract

We propose TWO-SIGMA-G, a competitive gene set test for scRNA-seq data. TWO-SIGMA-G uses a mixed-effects regression model based on our previously published TWO-SIGMA to test for differential expression at the gene-level. This regression-based model provides flexibility and rigor at the gene-level in (1) handling complex experimental designs, (2) accounting for the correlation between biological replicates, and (3) accommodating the distribution of scRNA-seq data to improve statistical inference. Moreover, TWO-SIGMA-G uses a novel approach to adjust for inter-gene-correlation (IGC) at the set-level to control the set-level false positive rate. Simulations demonstrate that TWO-SIGMA-G preserves type-I error and increases power in the presence of IGC compared to other methods. Application to two datasets identified HIV-associated Interferon pathways in xenograft mice and pathways associated with Alzheimer’s disease progression in humans.

## 1 Introduction

Single-cell RNA sequencing (scRNA-seq) data have been used to understand the heterogeneity of cell type landscapes and to understand the variation of gene expression at the single-cell resolution across biological processes or treatments. Many of the data analysis methods originally designed for bulk RNA-seq data, including Differential Expression (DE) based gene set tests, have been re-applied to scRNA-seq data. These DE-based gene set tests are used to test whether a pre-constructed [2] set of genes is significantly differentially expressed between/among sample groups. In the past decade and a half, such gene set tests have been used to contextualize gene-level differential expression analyses and identify both important pathways and real biological mechanisms [15, 21, 30, 14]. These gene set tests also improve statistical power and reduce spurious associations as compared to gene-level DE testing [8, 12]. This further increases reproducibility across multiple experiments, which is often lower than desired due to biological and technical variability across different transcription profiling platforms [8, 12]. Such DE-based gene set tests therefore constitute a routine step of performing differential expression (DE) based analyses, in both bulk RNA-seq and scRNA-seq datasets [31, 7].

There are at least two different types of DE-based analyses for scRNA-seq data. The first type of DE-based analysis focuses on the difference between sample groups to identify genes or gene sets significantly associated with sample groups. Example comparisons include disease vs control, mutant vs wild-type, and before vs after treatment. Questions of this type of analysis are similar to DE analysis in bulk RNA-seq data analysis. In scRNA-seq data, these analyses are typically performed within one cell type, meaning *p*-values of genes or gene sets are cell-type-specific. Another type of DE analysis is typically conducted by finding biomarker genes for a cell type, that is, to compare between or among cell types. This is often done by pooling samples in all sample groups together and assuming that gene expression differs primarily by cell type and not by sample group. In this paper we will focus on the first type of analysis, however the method we propose can perform gene set testing for both types of questions.

Conducting gene-level DE analysis is the first step in DE-based gene set testing. Therefore, it is important to use a method for gene-level DE analysis which employs suitable distributional assumptions to model scRNA-seq data and test for DE. One such method is our previously published TWO-SIGMA [38], which has several useful features. First, TWO-SIGMA assumes that observed counts follow a Zero-Inflated Negative Binomial (ZINB) distribution, which provides a flexible approach to capture the zero-inflated and overdispersed counts often seen in scRNA-seq, but not in bulk RNA-seq, datasets. Available DE-based gene set tests originally developed for microarray and bulk RNA-seq, including GSEA [34] and related extensions `sigPathway` and fGSEA [19], CAMERA [43], and PAGE [18], all conduct gene-level analyses using either non-parametric rank-based methods or using distributional assumptions chosen to represent the features of microarray or bulk RNA-seq data. Misspecified gene-level models can lead to misleading inference at both the gene-level and set-level, taking the form of inflated type-I error or reduced power. Second, TWO-SIGMA accounts for the fact that standard scRNA-seq experiments often collect many cells from the same biological sample [14, 15, 16, 5, 25]. This helps to meet a recent demand for methods that can account for sample-to-sample variation [16]. Most of the scRNA-seq studies published to date ignore potential sample effects by pooling all cells in one treatment group or condition together. Ignoring the possible correlation between different cells can increase the false positive rate [38]. To consider these sample effects, TWO-SIGMA can model sample ID as a fixed effect or a random effect. Because (1) we are usually not interested in estimating the individual sample effect but interested in the sample-group (e.g. treatment) effect and (2) and some cells in different treatment groups may come from the same donor, a model including sample as a random effect term can improve both interpretation and control of the false positive rate. TWO-SIGMA’s random effect modelling framework allows for efficient computation of within-sample correlation between cells in scRNA-seq data that has a large number of cells from the same biological sample.

It is essential to discriminate between the two types of null hypotheses, competitive and self-contained, used in gene set testing. Competitive gene set tests evaluate significance by comparing the evidence of differential expression of a gene set to the evidence in a reference set of genes [18, 36, 8, 28]. In contrast, self-contained tests are commonly used to compare the similarity of gene expression patterns in a gene set across different data sets [13, 42], not relative to other genes. Competitive tests use a battery of gene sets, for example from the Molecular Signatures Database [34, 20], to rank sets and analyze which are the most significantly associated with a given phenotype. The results of competitive tests are easier to interpret, and competitive tests are more common in the literature today [43, 11]. Competitive tests usually permute genes or use parametric assumptions to construct a distribution of these summary statistics under the null hypothesis, often assuming that genes are independent of each other [43, 42, 13, 10]. Previous studies have shown that competitive tests which assume independence often suffer from inflated type-I error [43, 13, 10]. This is because genes within a given gene set tend to have a positive IGC, even under the null hypothesis. Ignoring this IGC under-estimates the variance of set-level summary statistics and can dramatically inflate type-I error through inducing a typically positive correlation in the marginal gene-level statistics [3, 11, 43, 8]. Therefore, it is critical that any competitive gene set test adequately account for IGC to provide statistically rigorous set-level *p*-values.

We are aware of two existing methods explicitly created for competitive gene set testing using scRNA-seq data: iDEA [23] and an extension of MAST [10]. iDEA jointly conducts gene-level DE testing using zingeR [39] and uses a Bayesian approach to produce set-level *p*-values. iDEA does not adjust for IGC, however, and may not detect the scenario in which the same proportion of genes are significant in the test and reference set but the magnitude of the association differs. MAST fits a log-normal hurdle model at the gene-level and uses a Z-test with a computationally-intensive bootstrapping procedure that was not studied in great detail to produce set-level *p*-values. We introduce more of iDEA and MAST in the Methods section. Many other scRNA-seq-related gene set tests with different goals have been developed, including BAGSE [17], UniPath [4], PAGODA [9], SCENIC-AUcell [1]. BAGSE is not a competitive test and has similar hypothesis as GSEA [34], consisting of a hybrid of the self-contained and competitive null hypothesis [13, 6, 36], PAGODA looks for coordinated variation and is not DE based, UniPath uses single-cell ATAC-seq (scATAC-seq) data in addition to scRNA-seq data, and AUcell in the SCENIC workflow is based on regulon activity.

Overall, there is a need for methodological advancements to tailor gene set testing frameworks to scRNA-seq data, and a need to evaluate the ability of methods designed for bulk data to provide statistically valid results when applied to scRNA-seq data. This paper develops TWO-SIGMA-G, a set-level framework for DE testing in scRNA-seq data using the competitive null hypothesis. To test for DE at the gene-level, TWO-SIGMA-G uses our recently developed TWO-SIGMA [38], providing a flexible mixed-effects zero-inflated negative binomial regression model for good fit to the data at the gene-level. The use of a regression-based framework allows complex experiments including multiple covariates to be analyzed and provides many choices for gene-level statistics depending on the priority of the analysis. To avoid the inflated type-I error often caused by commonly existing positive IGC in a biological gene regulation pathway, IGC is estimated using an innovative residual-based approach and explicitly adjusted for at the set-level. We demonstrate TWO-SIGMA-G outperforms existing competitive gene set testing methods using extensive simulation scenarios. Application of TWO-SIGMA-G to an HIV-related humanized mouse scRNA-seq dataset and an Alzheimer’s Disease human brain scRNA-seq dataset reveal exciting biological findings. TWO-SIGMA-G is available at https://github.com/edvanburen/twosigma.

## 2 Materials and Methods

### 2.1 TWO-SIGMA for Gene-Level DE Statistics

TWO-SIGMA fits a mixed-effects zero-inflated negative binomial (ZINB) regression model and simultaneously models, for cell *j* of individual *i*, the probability of dropout *p*_*ij*_ and the negative binomial mean *µ*_*ij*_ as follows:

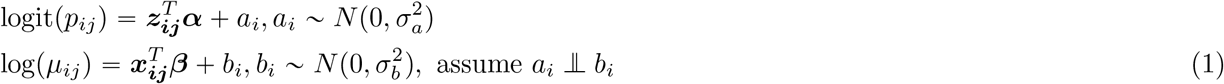

***α*** and ***β*** are fixed effect coefficient vectors and the corresponding vectors of covariates ***z***_***ij***_ and ***x***_***ij***_ can be different. *a*_*i*_ and *b*_*i*_ are sample-specific random intercept terms. Including these terms helps control for any within-sample correlation, providing more accurate estimates and standard errors of fixed effect parameters. The zero-inflation component can be removed in its entirety if desired, as is done in the real data analyses. In this context, gene-level statistics will correspond to tests of estimable contrasts of regression parameters from equation (1). Examples include a likelihood ratio statistic of a treatment effect, Stouffer’s method to combine the Z statistics [33, 10] from both components (if both are present), or an ANOVA style pairwise comparison between treatment groups within cell-type, as we demonstrate in the real data analysis of this paper. To summarize, TWO-SIGMA can control for additional covariates in both components, incorporate random effects to accommodate within-sample dependency, analyze unbalanced data, test general DE hypotheses beyond a two-group comparison, and allows for zero-inflated and overdispersed counts at the gene-level.

### 2.2 Estimation of Inter-Gene Correlation

Before specifying our new gene set testing method, we first propose a novel strategy to estimate IGC between pairs of genes from their respective gene-level DE regression models. Cell-level covariates such as the cellular detection rate (CDR), which measures the percentage of genes expressed in a cell, have been previously demonstrated to be highly influential to observed expression levels [10]. Subject-specific covariates, such as disease status or race, can further create an additional correlation structure in the raw data. Therefore, using the raw data to estimate IGC can overestimate the correlation that remains between gene-level statistics, which come from regression models that directly adjust for these other covariates. Thus, the use of residuals to estimate IGC can better represent the remaining correlation of the gene-level statistics under the null.

We estimate the inter-gene correlation of a given gene set using the residuals from the TWO-SIGMA model as follows: Define the (*n*_*i*_ ×1) vector of residuals for gene *s* from individual *i* as ***r***_***is***_ = ***Y***_***is***_ − ***Ŷ***_***is***_. Then, by individual, construct the *n*_*i*_ × *s* matrix ***R***_***i***_ = {***r***_***is***_} consisting of the residuals for all test set genes. Given these residual matrices, we can compute the pairwise (*s* × *s*) correlation matrix ***C***_*i*_, which contains *s* choose two unique non-diagonal elements. These elements give the pairwise correlations between the residuals of two different genes in the test set. We average these values to produce one average pairwise correlation 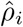 per individual. Finally, we estimate the overall correlation with the average of these values such that 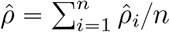

Our IGC procedure therefore builds off of the advantages of a residual-based approach in removing the correlation from sample-level and cell-level covariates. We further use individual-level calculations to help mitigate the impacts of the large individual heterogeneity often seen in scRNA-seq datasets. In simulations, we found that this IGC estimate preserves type-I error in a conservative manner while still producing improved power in a variety of realistic scenarios. The estimate of the IGC is virtually free computationally in that the model is not refit via permutation or bootstrapping.

Figure 2 shows the average set-level IGC estimates from TWO-SIGMA-G for each of the two comparisons in the Alzheimer’s dataset described more in the real data analysis section. A non-negligible correlation exists in the residual space for both comparisons, with half of the sets having a correlation larger than 0.02. For each comparison, over 98% of the estimated pairwise correlations are positive. Ignoring this remaining correlation would therefore make inflated type-I error a possibility.

### 2.3 TWO-SIGMA-G for Set-Level Testing

We extend our TWO-SIGMA method [38] to competitive gene set testing via TWO-SIGMA-Geneset (TWO-SIGMA-G), on overview of which is shown in Figure 1. Briefly, TWO-SIGMA uses a zero-inflated negative binomial regression model to test for DE at the gene level in scRNA-seq data. It is flexible and can be customized in several different ways. First, the zero-inflation component can be removed from the model entirely (as was done in the real data analysis section), leaving a standard negative binomial regression model. Second, the model can additionally include random effect terms to account for cell-cell correlation within the same sample and limit type-I error inflation in gene-level DE inference. Finally, many gene-level statistics measuring the evidence of DE can be used for set-level testing. For example, the likelihood ratio or z statistic corresponding to a treatment effect are commonly used gene-level statistics. We discuss uses for other, more complex, gene-level statistics based on custom contrasts of regression parameters in the real data analysis section. TWO-SIGMA-G employs the Wilcoxon rank-sum test to compare the statistics of genes in the test set to the statistics of genes in the reference set, and therefore uses the sum of the ranks in the test set as the set-level summary statistic. In using the ranks, TWO-SIGMA-G provides robustness against the influence of very large gene-level statistics.

**Figure 1:**
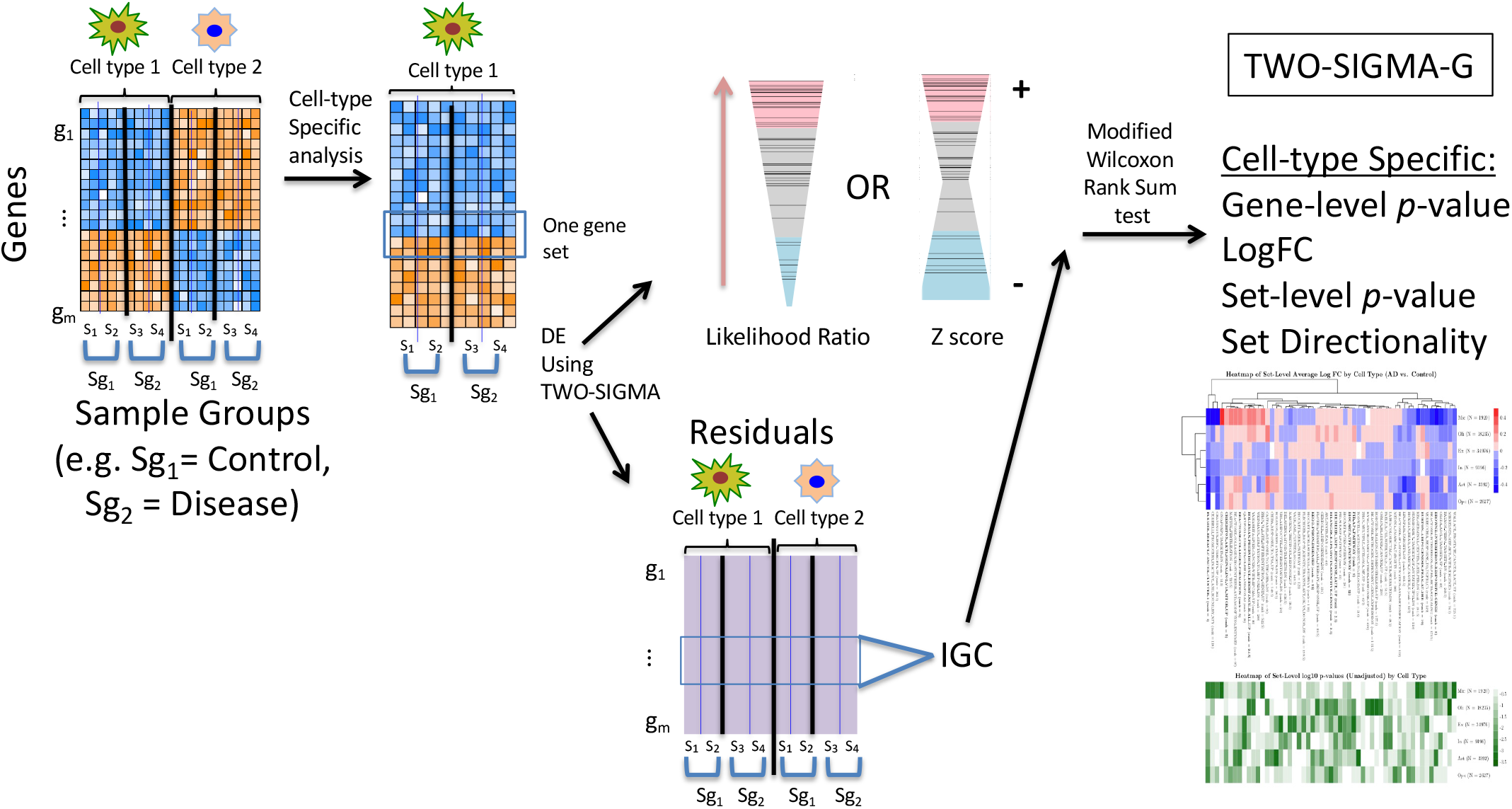
Overview of TWO-SIGMA-G. TWO-SIGMA-G enables cell-type-specific DE-based gene set testing using single-cell RNA-sequencing data. For clarity, we illustrate TWO-SIGMA-G on a binary comparison with two cell types, but we refer readers to the methods and real data analysis sections for examples of possible analyses involving more than two cell types and/or more than two sample groups simultaneously. In TWO-SIGMA-G, our previously published TWO-SIGMA method is used to produce gene-level statistics using the Likelihood Ratio test or the Z test. Inter-gene correlation (IGC) is estimated in a novel way using the gene-level residuals, and set-level inference is conducted using a modified Wilcoxon rank sum test which accounts for IGC to preserve type-I error. TWO-SIGMA-G is applicable both when analyzing DE between cell types for phenotype-associated gene sets or when analyzing DE between phenotypes for phenotype-associated gene sets.

**Figure 2:**
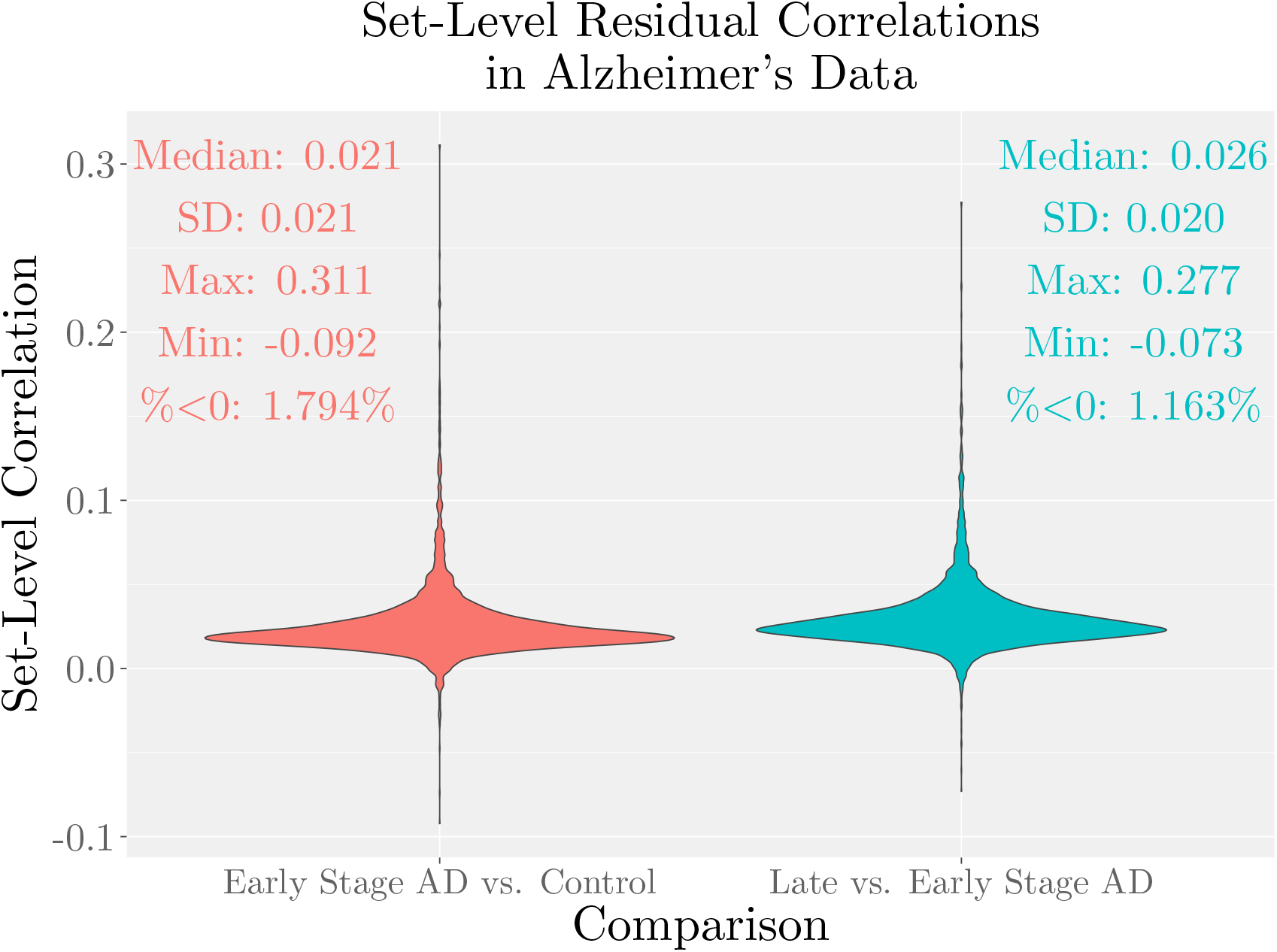
Set-level IGC estimates from TWO-SIGMA-G’s residual-based approach for the two Alzheimer’s dataset comparisons (see the real data analysis section). Sets plotted are taken from the c2 collection of the Molecular Signatures Database, and negative estimated correlations are set to zero to compute the TWO-SIGMA-G *p*-value in a conservative manner. Most sets demonstrate a substantially positive correlation after regressing out sample-level and cell-level covariates.

Traditionally, the Wilcoxon rank-sum test assumes that observations within a group are independent. However, as mentioned, IGC is expected given the construction of gene sets as harmonious biological pathways, and can inflate type-I error if ignored [43]. To create a gene set testing method designed for single-cell data, we therefore utilize a modified version of the rank-sum test. This modification allows for correlated gene-level statistics in the test set [3], similar to the approach of CAMERA for bulk RNA-seq [43]. We assume a pairwise correlation *ρ* between gene-level statistics in the test set of size *m*_1_ and no correlation in the reference set of size *m*_2_. With these assumptions, variance of the two-group Wilcoxon rank-sum statistic is:

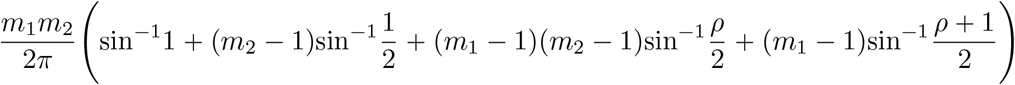

A positive *ρ* increases the variance as compared to a value of zero. Therefore, ignoring a positive *ρ* leads to an underestimated variance and inflated type-I error as a result. As discussed in the previous section, we estimate *ρ* using a residual-based approach. Using this modified variance, and the known mean of rank-sum statistics under the null, set-level *p*-values are computed analytically using a standard normal approximation [43]. The reference set used in TWO-SIGMA-G can be chosen in one of two ways: either using a random sample of other genes of size *m*_1_ or as the collection of all genes not in the test set under consideration.

In addition to producing set-level significance, TWO-SIGMA-G also identifies the directionality of sets as up or down-regulated. Whether or not a zero-inflation component is included in gene-level models, directionality is produced by averaging gene-level log fold-change estimates in the test set to produce a set-level effect size and taking the sign of the result. These effect sizes are demonstrated further in the real data analysis section.

As compared to other methods, TWO-SIGMA-G has several key advantages in applicability and interpretability. First, it is explicitly tailored to scRNA-seq data at the gene-level in that it can flexibly and optionally account for zero-inflation, overdispersion, and within-subject random effect terms to account for within-subject cell-cell correlation. Second, the use of a regression modeling framework at the gene-level enables the analysis of complex designs including multiple confounding covariates, as will be demonstrated further in the real data analysis section. Third, estimating IGC using residuals after regressing out sample-level and cell-level covariates provides estimates of IGC that more closely reflect the remaining correlation of the gene-level statistics.

### 2.4 Existing Methods

We compared TWO-SIGMA-G to six other methods for gene set testing: GSEA [34], fGSEA [19], PAGE [18], CAMERA [43], MAST [10], and iDEA [23]. Among them, PAGE, CAMERA, MAST, and our proposed TWO-SIGMA-G are competitive gene set tests. GSEA and fGSEA are both hybrid methods including components of both competitive and self-contained testing. For MAST and iDEA, we focused on their competitive testing procedures, and not their other functions, to be comparable to other five methods discussed in this paper.

GSEA [34], a very popular method, was developed as one of the original methods for gene set testing using bulk RNA-seq or microarray data. GSEA uses an enrichment score comparing two groups using a Kolmogorov-Smirnov-like test. The null hypothesis tested by GSEA is a hybrid of the competitive and self-contained null hypothesis. Larger gene sets are often more significant, even if additional genes represent noise, and other gene sets not being tested can influence results in counterintuitive ways [6, 36]. Although GSEA is not strictly a competitive test, we still include it for comparison given its popularity for performing gene set testing in both bulk and scRNA-seq data [31, 7]. GSEA was accessed via the `GSEA R` package (version 1.2, available at https://github.com/GSEA-MSigDB/GSEA_R). Other than the choice of 200 permutations, default options from the `GSEA R` package were used.

fGSEA was developed as an alternative to GSEA which relies on gene-level summary statistics to improve the computational performance of GSEA. Rather than estimating the null distribution of the enrichment statistic for each set separately, as is done in GSEA, fGSEA employs a custom procedure to simultaneously estimate the null distribution for all sets. was accessed via `fgsea R` package (version 1.10.1). Ten thousand permutations were used, and otherwise default options from the `fgsea R` package were used.

PAGE compares the log fold-change values in the test set to those from the complement set of genes using a one-sample z-test, where the sample mean and variance are estimated using all genes. PAGE assumes genes are independent and does not adjust for IGC, but is computationally efficient as compared to GSEA. It may fail to preserve type-I error in some cases, however. PAGE was accessed via the `PGSEA` function of the `PGSEA R` package (version 1.60).

CAMERA was developed as a competitive gene set test for microarray or RNA-seq data. Gene-level statistics are first constructed using a linear model, meaning that CAMERA can accommodate complex experimental designs beyond a two-group comparison. Set-level *p*-values are then computed using modifications of the t-test or Wilcoxon rank-sum test that allow for a common pairwise correlation in the test set. Rather than using the raw data to estimate the IGC, CAMERA uses the residuals from the linear model. As discussed above, use of the residuals means that the variation in gene expression explained by the covariates is removed, giving the most reliable estimate of the correlation between the gene-level statistics in the test set. By avoiding permutation, and unlike some early approaches, CAMERA provides a statistically valid, computationally efficient test of a precisely defined and fully specified null hypothesis [13]. The hypothesis corresponds to a test that the average absolute value of each coefficient in the test set is larger in magnitude as compared to the reference set. CAMERA was accessed via the `limma R` package (version 3.42).

MAST, which was developed for scRNA-seq DE analysis, has an extension to allow for competitive gene set testing comparing a test set to its complement set of genes [10]. This extension is quite flexible given the log-normal hurdle regression framework employed by MAST. Once the gene-level statistics are collected, a bootstrap procedure is used to estimate the inter-gene correlation of the regression coefficients for each component of the hurdle model. Set-level tests are conducted using the a modified Z-test which adjusts for the IGC and computed separately for the two components of the hurdle model. The performance of the method does not seem to have been studied in great depth, and recent evidence has suggested that log transforming scRNA-seq data may distort true signal [37, 22]. The gene set testing procedure of MAST was accessed via the `gseaAfterBoot` function of the `MAST R` package (version 1.13.5).

iDEA was developed as an integrative method for both DE and gene set enrichment analysis [23]. The method takes gene-level DE summary statistics and gene sets as input. For each gene set, the method produces a posterior probability of DE for each gene and a set-level *p*-value. As a competitive test, the set-level *p*-value compares the gene-level odds of DE in the test set to the reference set. Because it focuses on the posterior probability of DE at the gene level, iDEA may not capture a gene set where enrichment of the test and reference sets is similar in proportion but the effect sizes themselves are systematically larger in the test set. iDEA was accessed via the `iDEA R` package (version 1.0.1).

Both MAST and iDEA use the complement set of genes as the reference set. Previous studies have, however, cautioned that set size may inflate the type-I error of some gene set testing procedures [6, 36]. For larger gene sets, which are likely of more interest scientifically, the difference between these two approaches for choosing a reference set diminishes.

### 2.5 Simulation Studies

We utilize a custom simulation procedure to simulate correlated gene sets (See Supplementary Section S1 for full details) and test for set-level differential expression of a binary covariate representing a treatment effect. First, independent genes were simulated from the zero-inflated negative binomial distribution, optionally including (1) within-sample random effect terms to create a within-sample correlation structure, and (2) additional confounding covariates to create additional cell-cell correlation. For each independent gene, we then simulated 29 correlated genes by adding random noise from the negative binomial distribution to create correlated gene sets of size 30. Under the alternative, the magnitude of the added noise is increased to preserve signal and maintain the correlation structure. We simulated 1,000 independent genes without gene-level random effect terms and 300 independent genes with gene-level random effect terms for each of six settings (Supplementary Table S1) which vary the magnitude of added noise and the presence of additional covariates in the gene-level simulation. These six settings are intended to represent the diversity seen in real data sets to paint an accurate picture of testing properties over a wide range of gene sets and correlation structures. Simulations assumed 100 cells from each of 100 samples, and constructed uncorrelated reference sets randomly. To increase variability in the design matrix and thus the simulated expression data, we repeat this simulation procedure 10 times using the same distributional assumptions, but a different random seed. This simulation procedure not only allows us capture the impact of excess zeros, but it is particularly designed to vary the percentage of DE genes in the test and reference sets to mimic real data, which typically has has DE genes in both the test set and in the reference set. For example, the scenario “T50, R20” is a scenario under the competitive alternative hypothesis in which 50% of genes in the test set are DE and 20% of the genes in the test set are DE. Genes that are DE have the same effect size in all cases with the exception of the “mixed DE” scenarios (Supplementary Figure S8), which create gene sets with two different gene-level DE effect sizes.

We compared TWO-SIGMA-G to six popular methods for gene set testing. First, TWO-SIGMA-G was compared to three methods for gene set testing which utilize the full expression data: GSEA [34], CAMERA [43], and MAST [10]. These simulations were calibrated to produce gene-level statistics which summarize evidence from both the mean and zero-inflation components after adjustment for confounding covariates. Second, TWO-SIGMA-G was compared to three other highly relevant methods which rely on gene-level summary statistics: iDEA [23], PAGE [18], and fGSEA [19]. Simulations were conducted separately because the marginal gene-level effect sizes produced from informative scenarios evaluating the first three methods were modest, which led to difficulties obtaining reliable *p*-values from the second set of methods. Thus, to provide a fair and interesting comparison to the summary-statistic-based methods, we simulated correlated genes using the same framework, but increasing the signal in the mean component (Supplementary section S1.2). `R` version 3.6.3 was used to conduct simulations. The likelihood ratio test was used to calculate gene-level statistics in TWO-SIGMA-G, and default options were used for all methods unless otherwise specified. For fair comparison, log fold-change values from TWO-SIGMA were used as input for iDEA, PAGE, and fGSEA.

We primarily use boxplots to summarize simulation results. Each boxplot aggregates six different settings (Supplementary Table S1) which vary both the magnitude of the average inter-gene correlation (where applicable) in the test set and the nature of the correlation structure via the introduction of other individual-level covariates (figures 3, 4, and 5 and Supplementary Figures S3 to S8).

**Figure 3:**
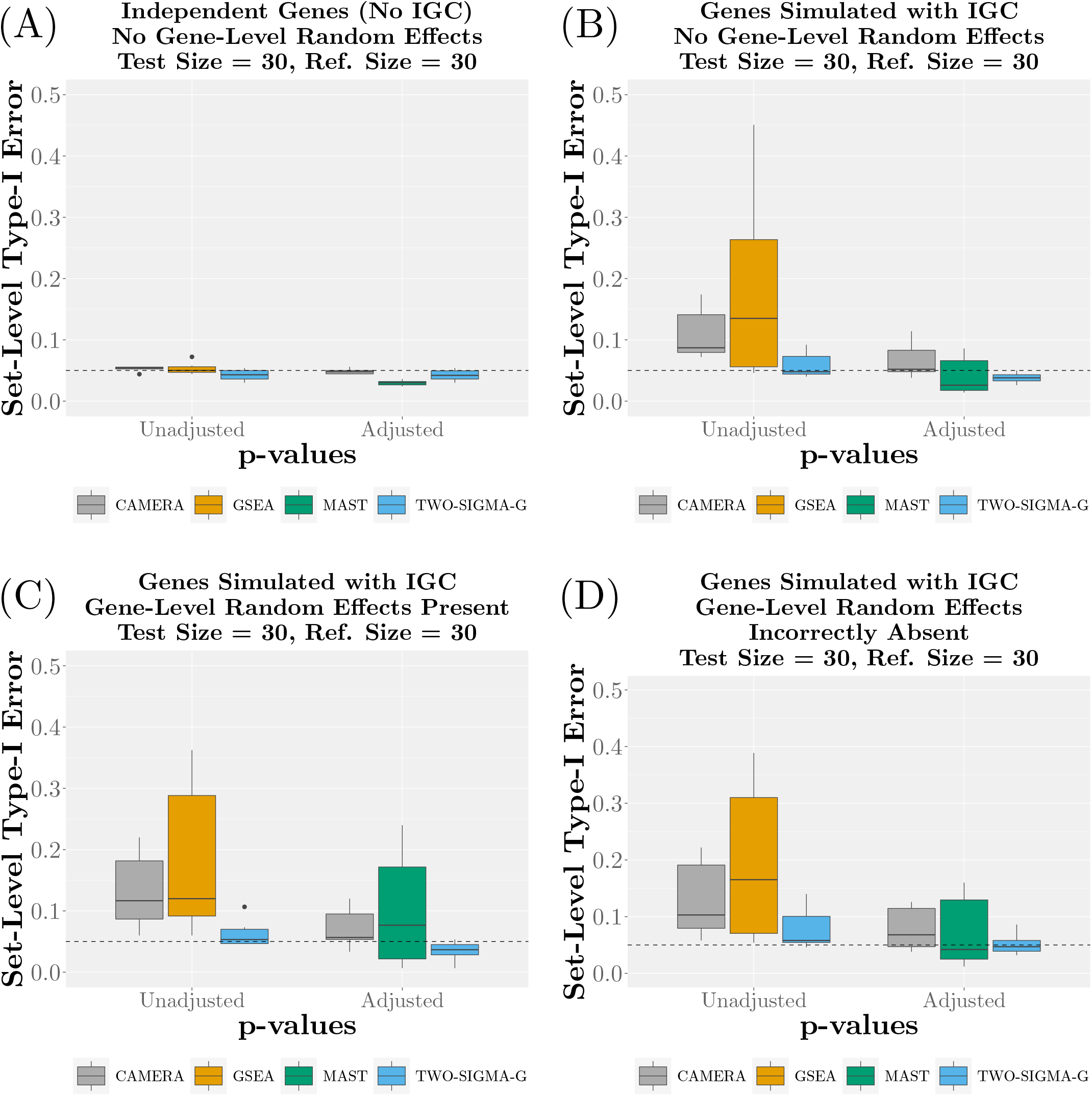
Type-I error performance for CAMERA, GSEA, MAST, and TWO-SIGMA-G using a reference set size of 30 genes. GSEA, CAMERA, and MAST were chosen for comparison here because all utilize the raw expression data, and the latter two use a regression modeling framework that can explicitly analyze complex experimental designs similar to the data analyzed in the real data analysis section. See Figure 4 for additional comparisons to summary-statistic based gene set testing methods fGSEA, iDEA, and PAGE. Each subfigure varies the existence of IGC between genes in the test set and the presence of gene-level random effect terms in the gene-level model (CAMERA and GSEA never include gene-level random effect terms). Within each subfigure, both unadjusted and adjusted set-level *p*-values are plotted, where available. See the Methods section of the main text and Supplementary Section S1 for more details regarding the simulation procedure. Significance was determined using a *p*-value threshold of 0.05, and 30 genes were included in the test and reference sets.

**Figure 4:**
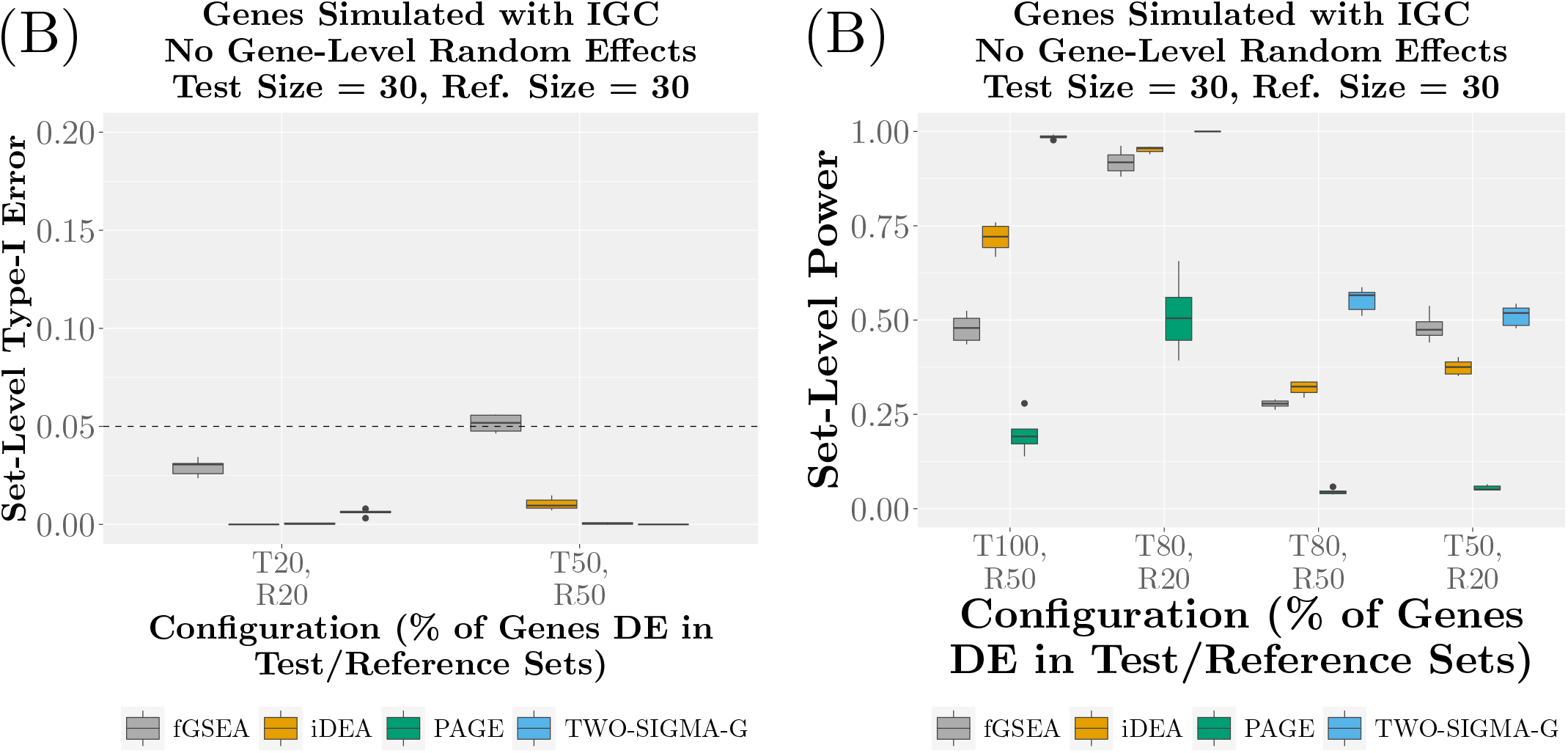
Type-I error (A) and power (B) performance of fGSEA, iDEA, PAGE, and TWO-SIGMA-G for various for genes simulated with IGC using a reference set size of 30 genes. fGSEA, iDEA, and PAGE are compared together because all use gene-level summary statistics instead of raw expression data. Scenarios along the *x*-axis vary the percentage of genes that are differentially expressed (with the same effect size) in the test and reference sets. Because of misleading model performance in cases with “R0,” these scenarios were excluded from the summary-statistic-based simulation studies. Significance was determined using a *p*-value threshold of 0.05, and 30 genes were included in the test and reference sets. See the Methods section of the main text and Supplementary Section S1 for more details regarding the simulation procedure.

**Figure 5:**
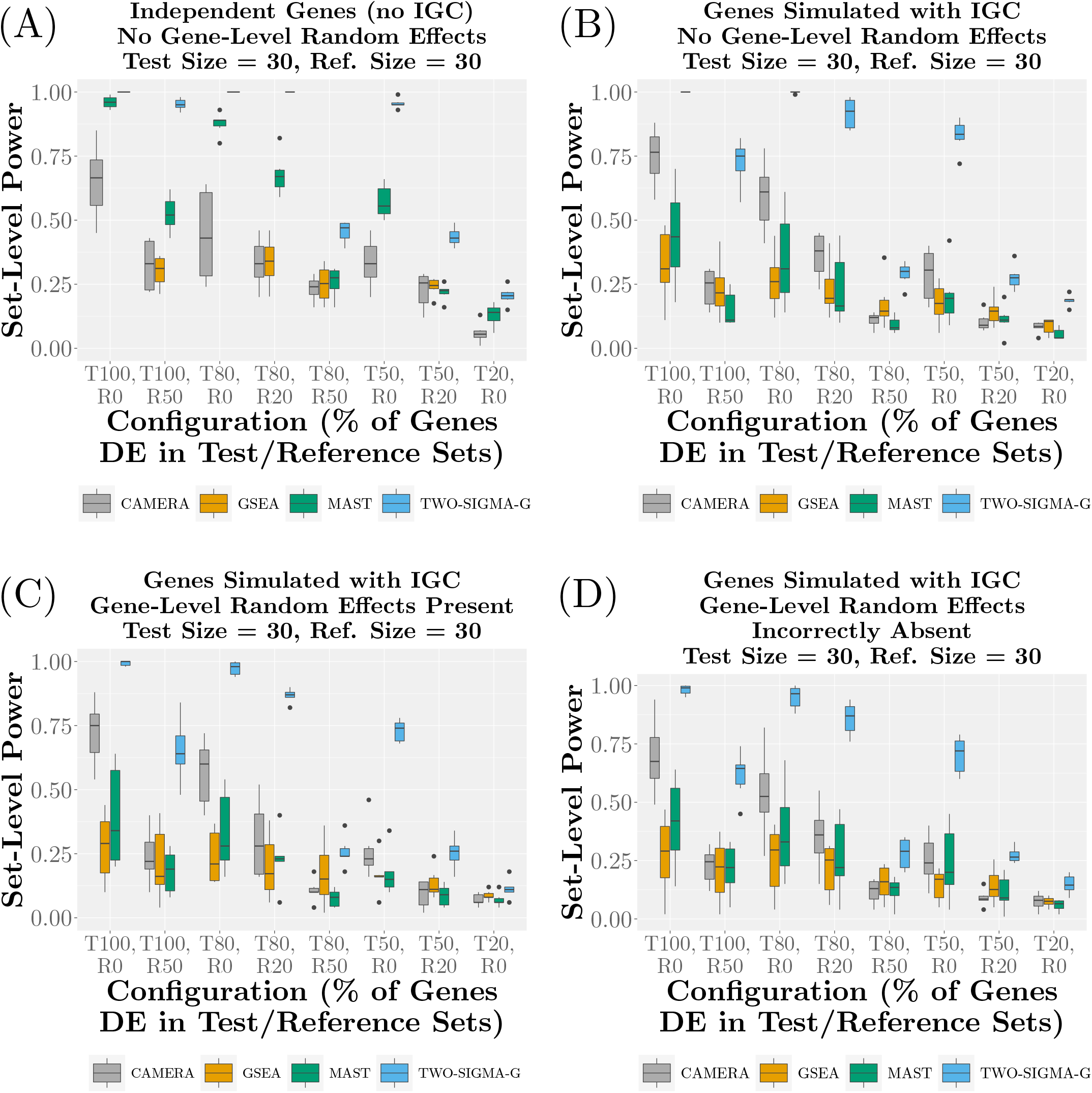
Set-level power of CAMERA, GSEA, MAST, and TWO-SIGMA-G using a reference set size of 30 genes. GSEA, CAMERA, and MAST were chosen for comparison here because all utilize the raw expression data, and the latter two use a regression modeling framework that can explicitly analyze complex experimental designs similar to the data analyzed in the real data analysis section. See Figure 4 for additional comparisons to summary-statistic based gene set testing methods fGSEA, iDEA, and PAGE. Each subfigure varies the existence of IGC between genes in the test set and the presence of gene-level random effect terms in the gene-level model (CAMERA and GSEA never include gene-level random effect terms). Scenarios along the *x*-axis of each subfigure vary the percentage of differentially expressed genes (with the same effect size) in the test and reference sets. For example, “T80,R50” corresponds to the configuration under the alternative hypothesis in which 80% of test set genes are DE and 50% of reference set genes are DE. Significance is determined using a *p*-value threshold of 0.05, and 30 genes were included in the test and reference sets (Supplementary Figure S7 presents results using a reference set of 100 genes). Note that GSEA did not output *p*-values for the “R0” scenarios. See the Methods section of the main text and Supplementary Section S1 for more details regarding the simulation procedure.

### 2.6 Two scRNA-seq data sets

We performed gene set testing on two different datasets to illustrate the usefulness of TWO-SIGMA-G. Gene sets were taken from the Molecular Signatures Database (mSigDB) [34, 20] version 7, c2 collection, accessed via the `msigdf` R package (https://github.com/ToledoEM/msigdf). All gene sets with at least two genes present after filtering were analyzed in each of two datasets described below. We used Fisher’s method to combine *p*-values over all cell types and create a consensus ranking of pathways (see, for example, Figure 6 (A)). Both datasets were from 10X UMI-based single cell RNA-seq platforms. As with other UMI-based scRNA-seq count data, we found that this data was not consistent with zero-inflation [35], and thus we fit the TWO-SIGMA model without the zero-inflation component at the gene-level.

**Figure 6:**
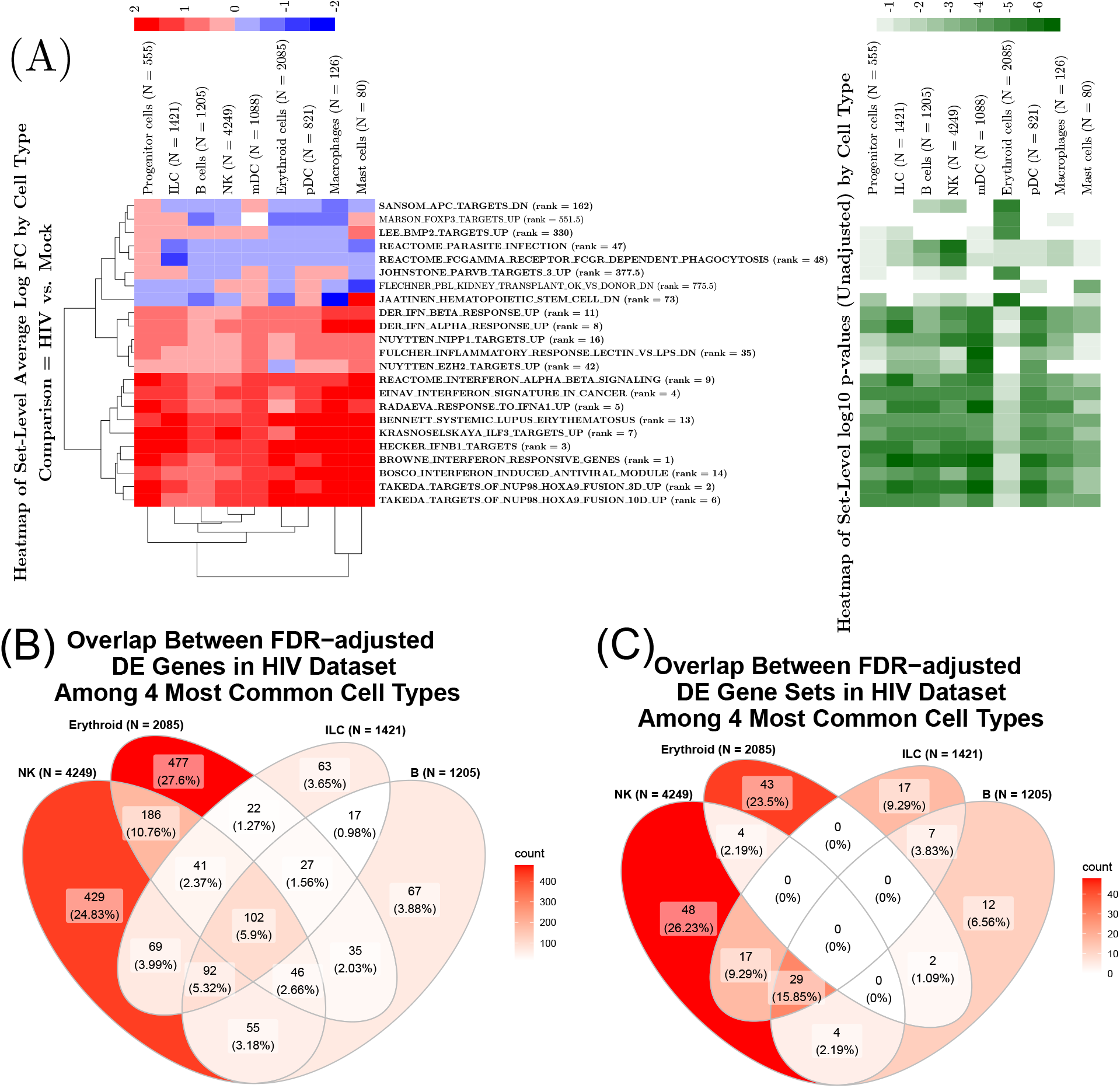
Results from analysis of the HIV dataset using TWO-SIGMA-G. (A) Cell-type specific variation in average set-level log fold-change (left) and significance (right). Sets plotted are among the top 10 in significance for at least one cell type. Sets in bold are significant at the 5% level over all cell types after FDR-adjustment of the Fisher’s method *p*-value, and the rank of the Fisher’s *p*-value among all sets is provided next to the set name. (B) Overlap between FDR-adjusted DE genes (5% significance level) among the four most prevalent cell types. (C) Overlap between FDR-adjusted DE gene sets among the four most prevalent cell types.

#### 2.6.1 HIV Dataset

Our first dataset consists of 11,630 single cells collected from humanized mice [5]. We kept only genes with a proportion of zeros no higher than the mean percentage, leaving the most relevant and highly expressed 3,549 genes. A total of nine cell types are present in the data: 4,249 natural killer (NK) cells, 2,085 erythroid cells, 1,421 innate lymphoid cells (ILC), 1,205 B cells, 1,088 myeloid dendritic cells (mDC), 821 plasmacytoid dendritic cells (pDC), 555 progenitor cells, 126 macrophage cells, and 80 mast cells. The read counts are then treated as the outcome of interest. Because the primary interest is in comparisons between HIV and mock cells within cell-type, we categorize cells into one of 2^*^9 = 18 mutually exclusive groups. An ANCOVA model additionally adjusting for CDR was fit as a way to test for cell-type specific differences in expression levels comparing HIV to mock. TWO-SIGMA-G is ideal for this analysis because gene-level statistics can come from a test of such an arbitrary contrast matrix. These gene-level statistics are, for each cell-type, Wald Z-statistics contrasting the mean values in observed expression between the two groups within a cell-type. After filtering to keep sets with at least two genes present in our data, a total of 4,772 sets from the MsigDB c2 collection were analyzed.

#### 2.6.2 Alzheimer’s Dataset

Our second dataset consists of 70,634 single cells from human donors [25]. A total of 48 individual donors are present, categorized into three pathology groups: 24 individuals are control patients free of a diagnosis of AD, 12 were diagnosed with early stage AD, and 12 were diagnosed with late stage AD. Cells from the six most common cell-types were analyzed: 34,976 excitatory neurons (Ex), 18,235 oligodendrocytes (Oli), 9,196 inhibitory neurons (In), 3,392 astrocytes (Ast), 2,627 oligodendrocyte progenitor cells (Opc) and 1,920 microglia cells. We did not remove cells beyond what was done in the original manuscript because extensive quality control was performed on the dataset we used. We chose to filter the original 17,926 genes to the 6,048 most highly expressed genes by removing genes unexpressed in at least 90% of cells. The read counts are once again treated as the outcome of interest in a model without a zero-inflation component. The existence of the pathology groups allows us to explore cell-type specific variability in gene expression as AD progresses into early and late stages of disease severity. Our geneset analysis was conducted similarly to above: a one-component ANCOVA model was fit including cell-type and AD status jointly, with age at death, sex, and the CDR used as an additional covariates. In total, 5,074 sets with at least two genes present from the MsigDB c2 collection were analyzed.

## 3 Results

### 3.1 Overview of TWO-SIGMA-G and Summary of Simulation Design

TWO-SIGMA-G is a gene set testing method based on differential expression specifically designed for scRNA-seq data. Because proper gene-level DE testing is key for a gene set test, TWO-SIGMA-G employs a mixed-effects zero-inflated negative binomial (ZINB) regression model we previously developed [38] to test for DE at the gene-level. This model can simultaneously capture the distribution of scRNA-seq datasets, account for within-sample correlation, analyze complex experiments by controlling for additional covariates, and provide options for gene-level test statistics. If zero-inflation exists at the gene-level, a gene-level model including both logistic and negative binomial components is fit. Otherwise, a one-component negative binomial model is used. When the ZINB model is used, gene-level DE test statistics can either (1) summarize the effects of both components using either the likelihood ratio test or Stouffer’s method [33, 10] to combine Z statistics or (2) be Z statistics testing a contrast of regression parameters in the mean component. For simplicity, random effect (RE) terms are included for all genes tested or for no genes. Then, the ranks of genes based on the gene-level DE test statistics will be further used in the gene set testing procedure. Specifically, set-level DE is tested in TWO-SIGMA-G using a modified version of the Wilcoxon rank sum test which adjusts for gene-gene correlation in the test set to control type-I error. As we will show below, TWO-SIGMA-G can improve power over other existing gene set testing methods that use either the gene expression matrix or gene-level summary statistics as input. The standard output of TWO-SIGMA-G includes gene-level DE summary statistics and associated *p*-values, set-level *p*-values and IGC estimates, and set-level effect sizes to characterize pathways as ‘Up’ or ‘Down’ regulated. When multiple cell types are included as a contrast, all information except IGC estimates is cell-type-specific, allowing users to investigate cellular heterogeneity.

We applied TWO-SIGMA-G to two scRNA-seq datasets, one experimental and one observational. TWO-SIGMA-G is freely available in the `twosigma R` package at https://github.com/edvanburen/twosigma. Full details of our proposed TWO-SIGMA-G is described in the Methods section.

We also performed extensive simulations to compare TWO-SIGMA-G to six other methods designed for gene set testing, the full details of which are given in Supplementary Section S1. Our real-data inspired simulation procedure is designed to reflect the zero-inflation, overdispersion, cell-cell correlation, and gene-gene correlation seen in counts using parameters inspired by real data and using six different settings (Supplementary Table S1). We simulated genes using a zero-inflated negative binomial distribution, optionally including random effect terms to simulate the presence of within-sample correlation. We then simulated correlated gene sets of size 30 using a custom approach detailed in Supplementary Section S1.2. The procedure is designed to study the impact of the following on set-level type-I error and power from TWO-SIGMA-G and six other methods: the magnitude and complexity of within-sample (cell-cell) correlation, the magnitude and complexity of gene-gene correlation, the amount and strength of gene-level DE, the presence of additional confounding covariates in the gene-level model, and the size of the reference set of genes.

### 3.2 TWO-SIGMA-G preserves type-I error in the presence of inter-gene correlation

In simulations, we compared the performance of TWO-SIGMA-G with other six gene set tests under the null hypothesis that there are no significant gene sets among sample groups. These methods are discussed above and are either competitive tests or have some flavor of competitive tests (GSEA and fGSEA).

First, we compared TWO-SIGMA-G to three methods (GSEA, CAMERA and MAST) which use the full scRNA-seq data matrix as input and have been popularly used in scRNA-seq data. In the comparisons, we particularly investigated the how IGC, gene-level random effects (RE), and both IGC and RE together in simulated data affect the false positive rate (Type I error) in these methods. Besides GSEA, the other three methods are designed to account for IGC. All four methods provide strong type-I error control when no genes are DE and genes are simulated independently and no gene-level within-sample random effects exist (Figure 3 (A), also see Supplementary Figures S1 and S2 for results using smaller significance thresholds after IGC adjustment). In contrast, type-I error is consistently inflated when IGC is present and ignored (Figure 3 (B)-(D), Unadjusted). This is particularly true for GSEA, which does not adjust for IGC or account other confounding covariates. When IGC is present and the estimated average IGC is used to adjust the gene set testing p-values in CAMERA and TWO-SIGMA-G, both methods hold Type I error correctly. While CAMERA and MAST still have some level of type-I error inflation after IGC adjustment (from expected 0.05 to max 0.1 or 0.15), TWO-SIGMA-G perfectly preserves type-I error at the 5% level in the presence of IGC (Figure 3 (B)-(D), Adjusted). Interestingly, TWO-SIGMA-G only has small level of type-I error inflation (3 (B)-(D), Unadjusted) even without IGC adjustment. The improved performance of TWO-SIGMA-G as compared to MAST, and CAMERA is likely due to a combination of factors that lead to a misspecified model for the features of scRNA-seq data in the latter two methods. First, CAMERA and MAST use a log transformation of the data, which may distort true signals, particularly in the presence of many zero counts [37, 22]. Second, unlike TWO-SIGMA-G and MAST, CAMERA does not separately model the excess zeros in the data and may underestimate parameters relating to mean expression as a result. The procedure used in TWO-SIGMA-G to estimate and adjust for IGC is well-calibrated and produces valid set-level inference.

We have also considered scenarios in which RE exist in the simulated data. Figure 3 (C) and (D) show that the type-I error from TWO-SIGMA-G is preserved or approximately preserved when gene-level random effect terms are truly present and either correctly included (“present”) or incorrectly excluded (“incorrectly absent”) from the fitted gene-level model. For CAMERA, GSEA, and MAST, however, type-I error tends to be inflated on average and the variance in the type-I error across the six settings tends to increase in the presence of gene-level random effects. For all methods, however, this type-I inflation is much lower in magnitude than can exist at the gene-level [38]. This highlights an advantage of competitive gene set testing: because it makes a relative comparison to a reference set of genes, it is partially robust to the consequences of a systematic, gene-level misspecification. The real data analysis section further shows a large agreement in set-level inference from TWO-SIGMA-G regardless of random effect inclusion in the gene-level model.

We additionally found that the null distributions of all three methods are nearly identical with a larger reference set size (Supplementary Figure S3). However, in the interest of being conservative, we will evaluate performance using results in which the test and reference sets are of equal size.

We also evaluated type-I error at various set-level null hypotheses, in which an equal but non-zero percentage of genes in the test and reference sets are DE (with the same gene-level effect size, see Supplementary Figure S4). For example, the scenario in which 20% of genes are DE in both the test and reference sets is one set-level competitive null hypothesis. Generally, all methods except GSEA become more conservative as the proportion of DE genes increases.

Second, we compared TWO-SIGMA-G to three other methods which use gene-level summary statistics: iDEA, fGSEA, and PAGE. To have a fair comparison among these methods and TWO-SIGMA-G, we used simulations with larger gene-level DE effect sizes to get gene-level summary statistics. iDEA, fGSEA, and PAGE produced reliable *p*-values when when excluding scenarios involving a completely null reference set (i.e. “R0”). Therefore, we only are able to evaluate type-I error at other various set-level null hypotheses, in which an equal but non-zero percentage of genes in the test and reference sets are DE. For example, the scenario in which 20% of genes are DE in both the test and reference sets is one set-level competitive null hypothesis. Figure 4 (A) shows that all four methods approximately control type-I error when evaluated at two set-level null scenarios (“T20,R20” and “T50,R50”). TWO-SIGMA-G, iDEA, and PAGE all tend to become more conservative as the total percentage of DE genes increases, while fGSEA becomes slightly more anti-conservative in this scenario. When using a larger reference set (Supplementary Figure S5 (A)) fGSEA fails to hold type-I error in scenarios where gene-level random effects were simulated and are included in the testing model (Supplementary Figure S6, (A) & (C)), and all methods hold type-I error. When random effects are incorrectly absent in the testing model (Supplementary Figure S6 (B) % (D)), type-I error is preserved by iDEA, PAGE, and TWO-SIGMA-G, but not by fGSEA with a reference set of size 100.

### 3.3 TWO-SIGMA-G improves power over alternative approaches

Figure 5 shows the power of CAMERA, GSEA, MAST, and TWO-SIGMA-G using simulated data, and demonstrate that TWO-SIGMA-G is consistently the most powerful method. Different configurations are presented, involving a differing proportion of DE genes (with the same effect size) in the test and reference set. For example, “T100,R50” corresponds to the configuration in which 100% of genes in the test set are DE and 50% of genes in the reference set are DE. Scenarios that include DE and non-DE genes in both the test and reference set are the most informative to study because it is unlikely in real data to have a completely null reference set and/or a completely alternative test set. Results from Figure 5 suggest that power depends primarily on the proportion difference in DE between the test and reference set and less on the precise composition of the test and reference sets. For example, the “T80,R50” and “T50,R20” configurations have the 30% more DE genes in the test set than the reference set, and similar power profiles for all methods within all four subfigures of Figure 5. We found that using a reference set size of 100 tends to improve power for all methods and particularly for TWO-SIGMA-G (Supplementary Figure S7). This power increase does not seem to be a consequence of an increase in type-I error (Supplementary Figure S3). This provides some evidence in favor of using a larger reference set in lieu of a balanced reference set.

The power of different testing methods has been further compared when gene-level random effect terms are truly non-zero and either correctly included (“present”) or incorrectly excluded (“incorrectly absent”) from the fitted gene-level model (Figure 5 (C) and (D)). In either case, power is only slightly reduced versus the cases without gene-level random effects seen in Figure 5 (A) and (B). Thus, if interested primarily in set-level inference, the increased computational cost from gene-level random effect terms may not be necessary for valid and powerful inference. However, any set-level power loss may be acceptable to prevent the massive type-I error inflation that has been shown to occur at the gene-level when random effects are mistakenly absent if gene-level inference is of interest [38].

When the magnitude of gene-level DE is varied, such that half of genes have twice the effect size of the other half, we found that set-level power is improved (Supplementary Figure S8). The relative positions of each configuration remain as in Figure 5, however, suggesting that power results in Figure 5 apply to alternative DE breakdowns. For example, whether or not genes in the test have varying DE magnitudes, the “T80, R20” scenarios have improved power over the “T100,R50” scenarios. The relative rankings of the three compared methods also remains when the magnitude of gene-level DE is varied.

As above, we also used simulations with larger gene-level DE effect sizes to compare the power of TWO-SIGMA-G to iDEA, fGSEA, and PAGE. Broadly, all of the results discussed above also apply to the comparisons of the summary statistic based methods. Figure 4 (B) shows power for scenarios that do not involve a completely null reference set (i.e., without “R0”). TWO-SIGMA-G is the most powerful method across a variety of set-level alternative scenarios as compared to summary statistic based methods, however the magnitude of the difference depends on the precise configuration of genes in the test and reference sets. When using a reference set of size 100 (Supplementary Figure S9), power is uniformly improved for all methods, but the relative ranks of the methods remain the same. In scenarios where gene-level random effects exist (Supplementary Figure S10), all methods tend to have reduced power but remain in a similar relative ranking. One exception is seen supplementary Figure S10 (A), which shows that fGSEA is more powerful than TWO-SIGMA-G in the “T50, R20” scenario. Comparing Supplementary Figure S10 (A) and (B) to (C) and (D), respectively, shows that use of a larger reference set tends to increase power for all methods.

### 3.4 Analysis of HIV data reveals biologically expected findings

We analyzed a dataset of 11,630 single-cells collected from of 4 humanized donor mice, two of which were infected with HIV and two were given a mock treatment [5] (also described in the Methods section). A total of 3,549 genes and 4,772 gene sets from the Molecular Signatures Database (MSigDB) [34, 20] were analyzed. The number of differentially expressed genes and sets in all cell types comparing HIV to a mock treatment is shown in Table 1. An increase in the number of significantly DE genes does not always correspond to more significantly DE gene sets. For example, erythroid cells have the second largest number of DE genes, but rank eighth in terms of the number of DE gene sets. This observation is expected in a competitive gene set test like TWO-SIGMA-G, because competitive tests focus on the relative signal of gene sets as compared to a background reference set of genes. The lack of a direct relationship between the number of DE genes and gene sets is also reflected when analyzing the overlap in significance between genes and gene sets among the four most prevalent cell types as seen in of Figure 6 (B) and (C). Instead of being a drawback, this highlights the need of using gene set testing to give extra set-level information beyond that from individual-gene-level DE tests.

**Table 1:**
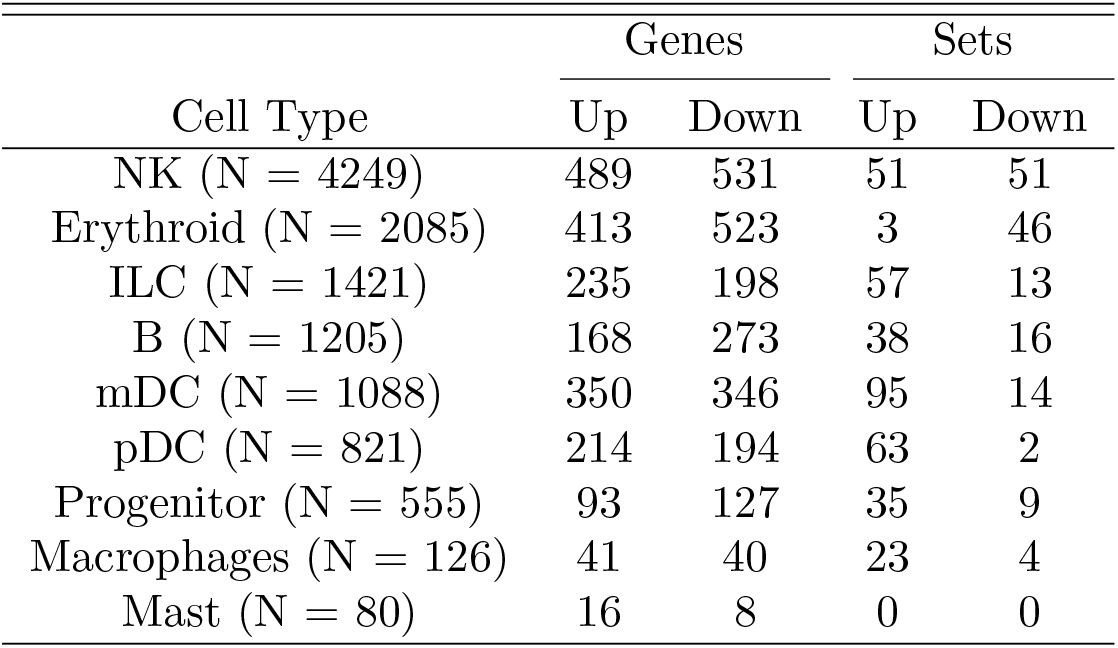
Shows the number of differentially expressed genes (using TWO-SIGMA) and gene sets (using TWO-SIGMA-G) after FDR-adjustment for the HIV dataset. Gene-level *p*-values were adjusted using the Benjamini-Hochberg method, and significance was determined by comparing these adjusted *p*-values to the 0.05 significance threshold. At the set-level, FDR-adjusted *p*-values were compared to the 0.2 threshold to mimic an exploratory analysis.

| Cell Type | Genes |  | Sets |  |
| --- | --- | --- | --- | --- |
|  | Up | Down | Up | Down |
| NK (N = 4249) | 489 | 531 | 51 | 51 |
| Erythroid (N = 2085) | 413 | 523 | 3 | 46 |
| ILC (N = 1421) | 235 | 198 | 57 | 13 |
| B (N = 1205) | 168 | 273 | 38 | 16 |
| mDC (N = 1088) | 350 | 346 | 95 | 14 |
| pDC (N = 821) | 214 | 194 | 63 | 2 |
| Progenitor (N = 555) | 93 | 127 | 35 | 9 |
| Macrophages (N = 126) | 41 | 40 | 23 | 4 |
| Mast (N = 80) | 16 | 8 | 0 | 0 |

Within-cell-type (”cell-type specific”) TWO-SIGMA-G results comparing HIV to the mock treatment are presented in Figure 6 (A) as cell-type-specific average log fold changes (FC) and corresponding *p*-values. In most cases, the main cell types share significance in common pathways associated with HIV infection, while in other cases there are HIV associated pathways which are significant only in a small number of cell types. In the gene sets among the ten most significant sets in at least one of the nine cell-types (Figure 6 (A)), sets related to virus introduction and interferon release are expected to be consistently upregulated and highly significant at both the set-level (as seen in a representative gene set in Supplementary Figure S11) and the gene-level [32]. The significance of these sets is found both when combining *p*-values into a consensus FDR-adjusted *p*-value using Fisher’s method and within cell types other than erythroid cells, albeit with differing strength of significance. Given the known functionality of erythroid cells as oxygen carriers in contrast to the immunological function of the other cell types, this result is expected. It also demonstrates that TWO-SIGMA-G can recover expected biological findings using cell-type-specific analyses and quantify differing strengths of association even among sets that may not exhibit large cell-type-specific heterogeneity. Figure 6 (B)-(C) show the overlap in significant DE genes and gene sets, respectively, after FDR-adjustment for multiple testing among the four most prevalent cell types. For example, Figure 6 (C) shows that there are 48 gene sets that are significant only in NK cells after FDR adjustment. These Venn diagrams show that our analysis reveals a large degree of cell-type specific heterogeneity at the gene level and the set level.

We also made comparisons between TWO-SIGMA-G, MAST, and CAMERA and briefly discuss their performance in NK cells (the most common cell type). We focused on comparisons to MAST and CAMERA because both can accommodate complex experiments via regression models and both estimate and adjust for IGC (see Figure 2 for IGC estimates from our other real data application) using the entire data matrix. Figure 7 (A) shows that only a moderate amount of overlap exists across the methods, with 16 sets shared among the top 50 in each of the three methods as ranked by *p*-value. Among these top 50 sets, TWO-SIGMA-G and CAMERA have a large overlap with 41 sets shared. MAST has much lower overlap with both TWO-SIGMA-G (18 sets) and CAMERA (17 sets). Although the sets “MOSERLE IFNA RESPONSE” and “BOSCO INTERFERON INDUCED ANTIVIRAL MODULE” are significant in MAST, they are not among the top 50 sets for MAST as they are for both TWO-SIGMA-G and CAMERA. This is unexpected given the nature of NK cells in response to HIV exposure [29, 26]. MAST has an extremely skewed *p*-value distribution (Supplementary Figure S13) and rejected 1,713 sets in NK cells after FDR adjustment (see Supplementary Figure S12 for before FDR-adjustment results), as compared to 102 sets for TWO-SIGMA-G and 188 for CAMERA (Figure 7 (B)). Although we did not explore the real data performance of MAST further, it is surprising that over 1,700 sets would be relevant to HIV introduction in only one cell type. Overall, we suspect MAST generated more false positives in real data as seen by the large number of significant gene sets and its reduced overlap with the other two methods among the top 50. In contrast, TWO-SIGMA-G and CAMERA perform similarly to identify significant interferon pathways in the HIV data. We also found similar patterns among other prevalent cell types (data not shown).

**Figure 7:**
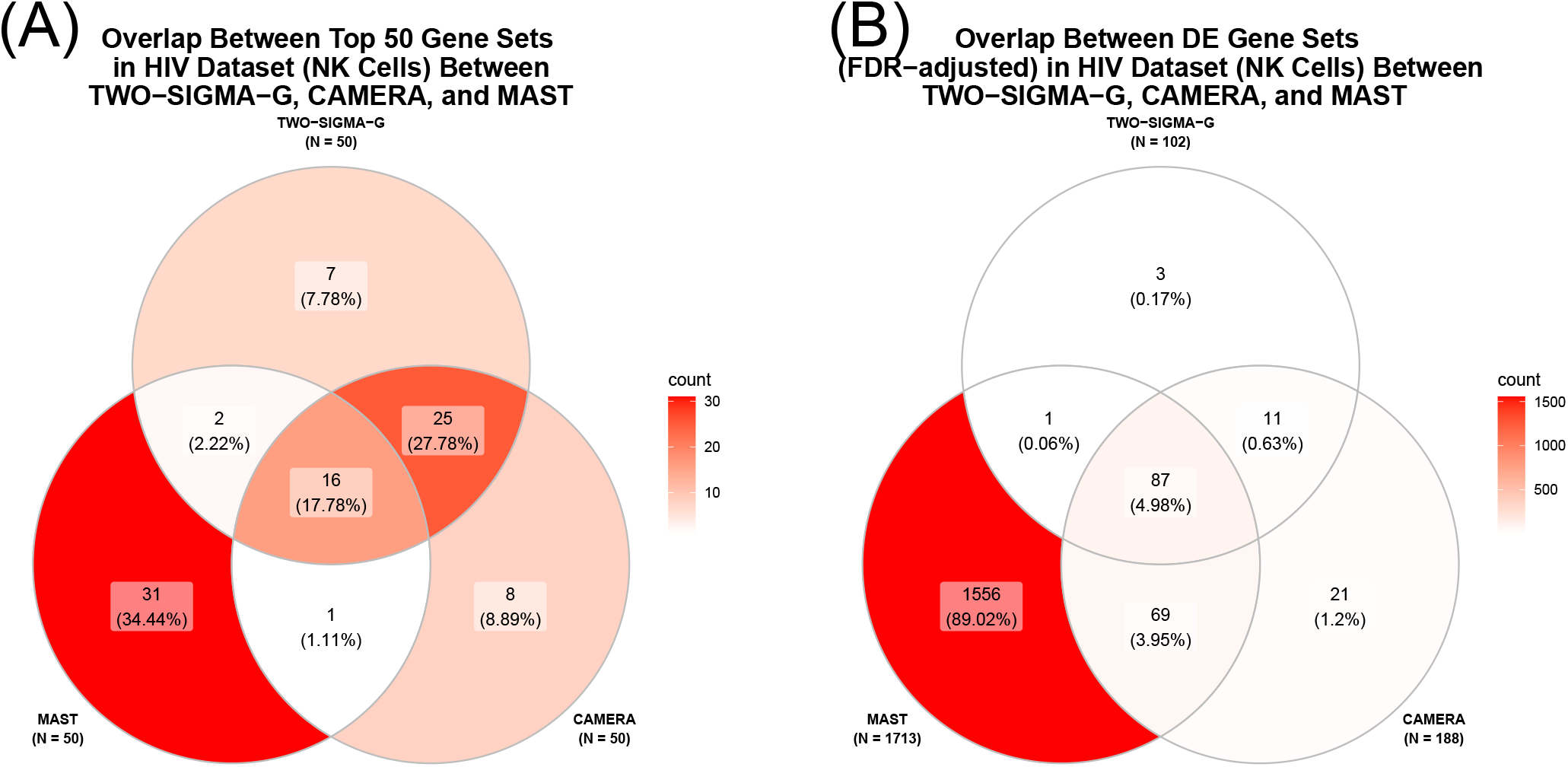
Comparison of the performance of TWO-SIGMA-G, CAMERA, and MAST in the HIV dataset. (A) Overlap among the top 50 gene sets for each method. (B) Overlap among all FDR-adjusted DE gene sets in NK cells.

### 3.5 Analysis of Alzheimer’s data reveals cell-type-specific heterogeneity in set-level expression

We chose to use a second scRNA-seq dataset to demonstrate how TWO-SIGMA-G can handle a more complex application. Specifically, we use the scRNA-seq data of [25] (see the Methods section for more details) to analyze changes in gene expression as Alzheimer’s Disease (AD) progresses. The data provides gene expression across three distinct pathology groups: control (AD free), early stage AD progression, and late stage AD progression. We focus on two relevant comparisons: late vs. early stage AD, and early stage AD vs. control. A total of 6,048 genes and 5,074 gene sets from the Molecular Signatures Database (mSigDB) [34, 20] were analyzed.

We used TWO-SIGMA-G to compare early stage AD patients to control in each of the cell types and provide cell-type-specific results (Figure 8). Previous studies have suggested that disruptions in mitochondrial functioning, particularly in cellular respiration as caused by oxidative damage, is among the earlist events in Alzheimer’s disease [27]. As Figure 8 (A) demonstrates, we replicate this finding with particularly robust downregulation seen in pathways related to cellular respiration, such as “KEGG OXIDATIVE PHOSPHORYLATION” and “MOOTHA VOXPHOS” (see Supplementary Figures S14 and S15 for more detailed gene-level results for these sets). The neuronal cell types demonstrate highly consistent statistical significance in these pathways. The Venn diagrams in Figure 8 (B) and (C) show that, after FDR correction, the majority of the overlap in significant genes and gene sets comes from the two neuronal cell types. Pathway changes in the early stages of Alzheimer’s disease therefore seem to manifest themselves as changes in the functioning of these two neuronal cell types. This result has been demonstrated previously [40].

**Figure 8:**
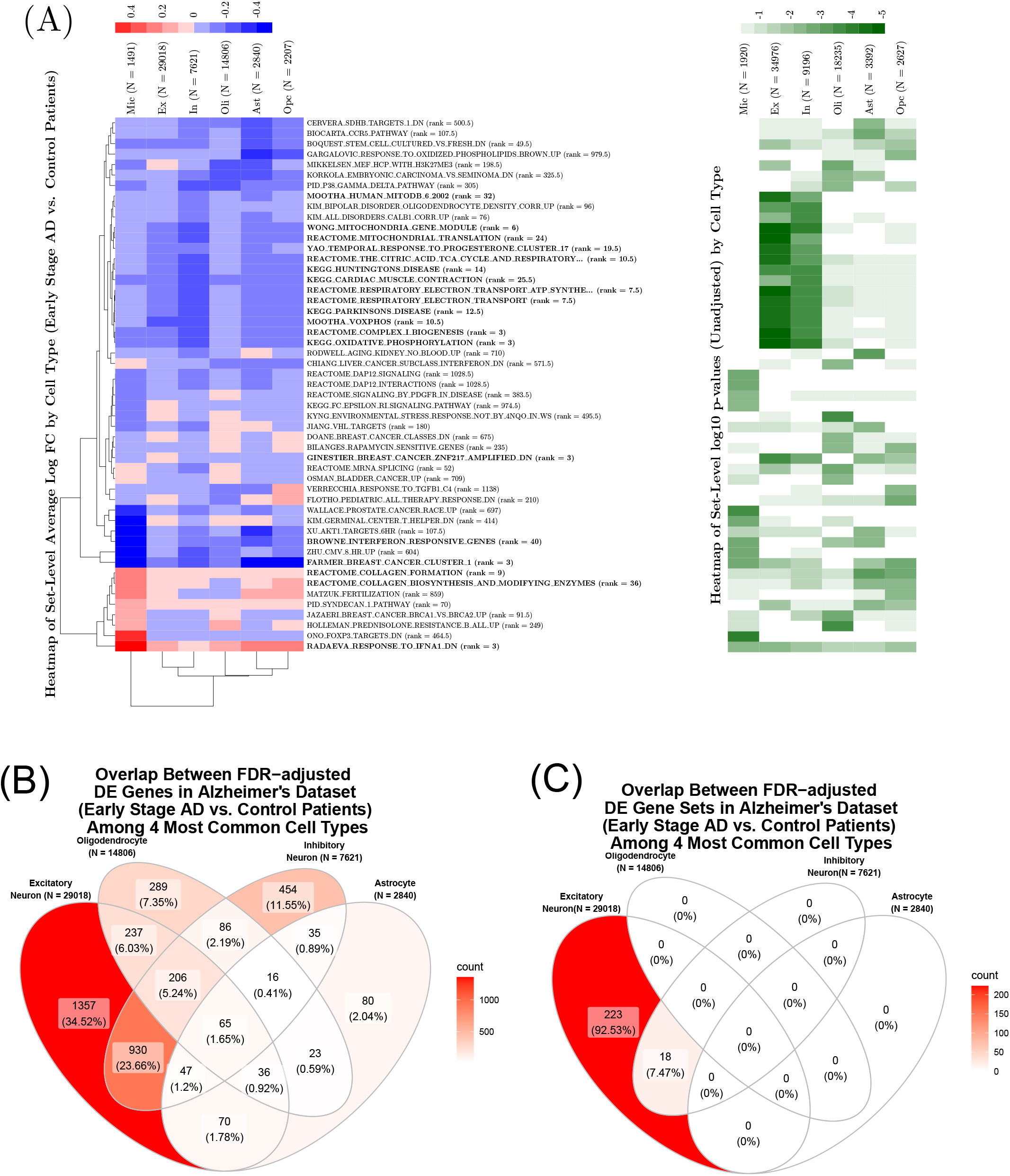
Results from analysis of Alzheimer’s dataset comparing Early Stage AD to Control using TWO-SIGMA-G. (A) Cell-type-specific variation in average set-level log fold-change (left) and significance (right). Gene sets plotted are among the top 10 in significance for at least one cell type. Sets in bold are significant at the 5% level over all cell types after FDR-adjustment of the Fisher’s method *p*-value, and the rank of the Fisher’s *p*-value among all sets is provided next to the set name. (B) Overlap between FDR-adjusted DE genes (5% significance level) among the four most prevalent cell types. (C) Overlap between FDR-adjusted DE gene sets among the four most prevalent cell types.

Previous differential expression analyses in AD patients have further suggested that this trend of decreased expression in genes associated with cellular respiration may reverse as the disease progresses [27, 24]. To investigate this possibility, Figure 9 shows cell-type-specific results comparing late stage AD to early stage AD. Heatmaps in Figure 9 (A) show that most of the top gene sets are now consistently and significantly upregulated, and furthermore many of these same sets are seen as highly significant but downregulated in Figure 8 (A). Thus, the initial downregulation in gene sets related to cellular respiration is reversed as AD progresses, possibly due to cellular degeneration and an increasing demand for energy in remaining cells [27] (see Supplementary figures S16 and S17 for gene-level information for the sets discussed above). Interestingly, this observed upregulation and the possible increase in demand for energy is highly significant in the neuronal cells, as in the previous comparison, but also highly significant in astrocytes, oligodendrocytes, and oligodendrocyte progenitor cells. When comparing late-stage AD to control (Supplementary figures S18-S20), there is a slight upregulation in respiration related gene sets, although such sets are not among the most significant sets for any cell type. Thus, our analyses suggest that there is a systematic decrease in the expression of genes related to cellular respiration in the neuronal cells of early-stage AD patients. This decrease is reversed and even slightly over-compensated for when comparing late-stage patients to early stage patients, both in neuronal cells and in other cell types. Without the breakdowns of AD patients into the early and late stages, this pattern is obscured (Supplementary figures S21-S23).

**Figure 9:**
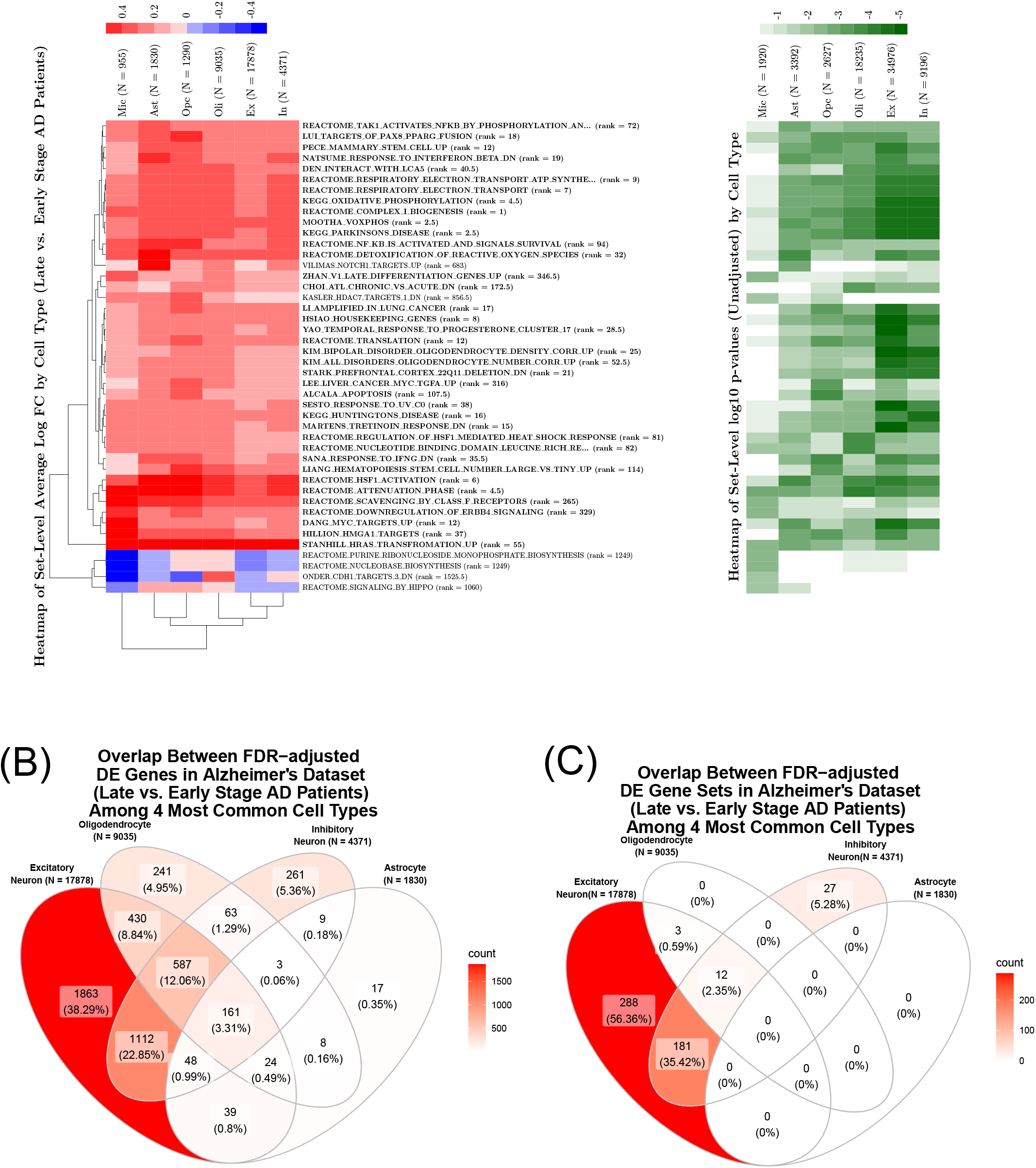
Results from analysis of the Alzheimer’s dataset comparing Late to Early Stage AD using TWO-SIGMA-G. (A) Cell-type-specific variation in average set-level log fold-change (left) and significance (right). Gene sets plotted are among the top 10 in significance for at least one cell type. Sets in bold are significant at the 5% level over all cell types after FDR-adjustment of the Fisher’s method *p*-value, and the rank of the Fisher’s *p*-value among all sets is provided next to the set name. (B) Overlap between FDR-adjusted DE genes (5% significance level) among the four most prevalent cell types. (C) Overlap between FDR-adjusted DE gene sets among the four most prevalent cell types.

The numbers of differentially expressed gene sets and genes for both comparisons in each of the cell types are summarized (Table 2). As mentioned above, most of the significant gene sets are downregulated when comparing early stage AD to control, but upregulated when comparing late to early stage AD. The number of significant genes and gene sets increases dramatically in the late vs. early-stage AD comparison over the early stage AD to control comparison. The fact that so many genes in the top three rows of table 2 are significantly DE reveals one reason that competitive set-level inference is useful in some scRNA-seq analyses. The large sample sizes of cells means that very modest differences in gene-level expression can be statistically significant, and thus that gene-level *p*-values must be interpreted cautiously. Competitive set-level analyses can contextualize gene-level differences and rank pathways to help highlight meaningful changes in important biological processes.

**Table 2:** Shows the number of differentially expressed genes (using TWO-SIGMA) and gene sets (using TWO-SIGMA-G) after FDR-adjustment for both comparisons of the Alzheimer’s dataset. Gene-level *p*-values were adjusted using the Benjamini-Hochberg method, and significance was determined by comparing these adjusted *p*-values to the 0.05 significance threshold. At the set-level, FDR-adjusted *p*-values were compared to the 0.2 threshold to mimic an exploratory analysis.

| Cell Type | Early Stage AD vs. Control |  |  |  | Late vs. Early Stage AD |  |  |  |
| --- | --- | --- | --- | --- | --- | --- | --- | --- |
|  | Genes |  | Sets |  | Genes |  | Sets |  |
|  | Up | Down | Up | Down | Up | Down | Up | Down |
| Excitatory Neuron (N = 29018, 17878) | 1055 | 1893 | 18 | 223 | 1619 | 2645 | 467 | 17 |
| Oligodendrocyte (N = 14806, 9035) | 339 | 619 | 0 | 0 | 1443 | 74 | 15 | 0 |
| Inhibitory Neuron (N = 7621, 4371) | 58 | 1781 | 0 | 18 | 1483 | 761 | 208 | 12 |
| Astrocyte (N = 2840, 1830) | 61 | 311 | 0 | 0 | 265 | 44 | 0 | 0 |
| Oligodendrocyte Progenitor (N = 2207, 1290) | 13 | 64 | 0 | 0 | 256 | 3 | 0 | 0 |
| Microglia (N = 1491, 955) | 27 | 20 | 0 | 0 | 14 | 0 | 0 | 0 |

Our analysis also reveals other cell-type type-specific heterogeneity. For example, microglia cells tend to have a unique set-level effect size profile, as demonstrated by the hierarchical clustering in the left heatmap of Figure 8. This uniqueness also extends to significance. In comparing early-stage patients to control, microglia cells exhibit stronger significance in pathways involved in immune response, such as “RADAEVA RESPONSE TO IFNA1 DN” or “BROWNE INTERFERON RESPONSIVE GENES,” while showing less or no significance in previously mentioned pathways related to cellular respiration. Given the role of microglia cells in immune response [44], these results are not surprising. For a general application, however, TWO-SIGMA-G can help researchers to investigate cell-type-specific heterogeneity when cellular functions are not understood. The ability to test complex gene-level hypotheses as contrasts of regression parameters increases the diversity of cell-type-specific hypotheses that can be explored.

Finally, we also compared the performance of TWO-SIGMA-G to MAST and CAMERA for both comparisons using the Alzheimer’s data (Supplementary Figures S24 to S27) in Ex cells (the most common cell type). Once again, we focused on comparisons to MAST and CAMERA because both can use regression models to control for additional covariates in this observational dataset and both estimate and adjust for IGC, which we found to be distinctly positive in this dataset (Figure 2). Figure 10 (A) and (C) show the overlap among the top 50 sets for the three methods. Each method identifies at least 10 sets uniquely for each of the two comparisons. As one example, when comparing late stage to early stage AD in Ex cells (Figure 10 (C)), TWO-SIGMA-G ranks 14 sets uniquely among its top 50, including the sets “MOOTHA VOXPHOS,” “KEGG OXIDATIVE PHOSPHORYLATION,” and “REAC-TOME RESPIRATORY ELECTRON TRANSPORT.” All methods identify these sets and their constituent genes as significant (see Supplementary Figure S28 for gene level results for the latter), but it is surprising that they are not identified among the 50 most significant sets for Ex cells given the known degradation seen in neuronal cells as AD progresses [41, 24].

**Figure 10:**
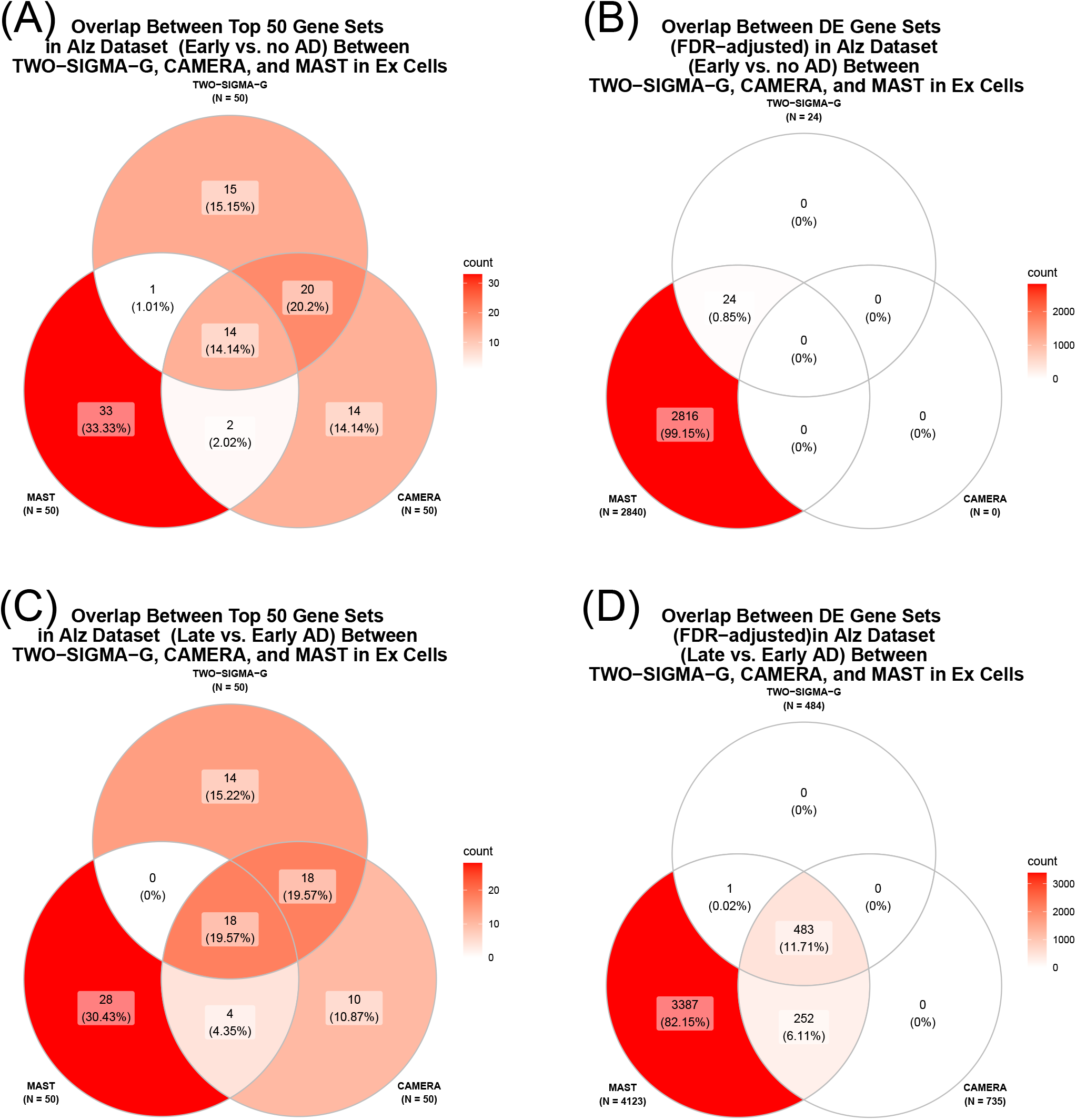
Comparison of the performance of TWO-SIGMA-G, CAMERA, and MAST in the Alzheimer’s dataset. (A) Overlap among the top 50 gene sets for each method in Ex cells comparing early stage AD to control. (B) Overlap among all FDR-adjusted DE gene sets in Ex cells comparing early stage AD to control. (C) Overlap among the top 50 gene sets for each method in Ex cells comparing late to early stage AD. (D) Overlap among all FDR-adjusted DE gene sets in Ex cells comparing late to early stage AD.

We find MAST rejects far more sets than both TWO-SIGMA-G and CAMERA after FDR-adjustment ((B) and (D) in Figure 10). The numbers of DE sets before FDR adjustment are seen in Supplementary Figures S24 and S26. Both Alzheimer’s comparisons also exhibit an extremely skewed *p*-value distribution in MAST (Supplementary Figures S25 and S27). Even after FDR adjustment, in the late vs. early stage AD and early stage AD vs. control comparisons, respectively, MAST rejected as much as 4,123 and 2,840 of a total 5,074 sets in Ex cells. Overall, as with the HIV data, we suspect that MAST is generating many false positives, but have not explored the reason for the massive number of rejected sets in more detail.

## 4 Discussion

We propose TWO-SIGMA-G, a novel method designed for competitive gene set testing using scRNA-seq data. At the gene-level, we employ our previously developed TWO-SIGMA method to test for DE. TWO-SIGMA is a flexible regression modelling framework that can fit both one-component and two-component negative binomial regression models to allow for overdispersed and zero-inflated counts. Additional covariates can be included in each of the two components, and sample-specific random effect terms can be included to account for within-sample correlation. The gene-level hypothesis is not limited to a binary or categorical group comparison, but rather can be a general contrast of regression parameters, as demonstrated in the real data analyses. This flexibility allows the testing of complex hypotheses and the analysis of complex experimental designs. At the set-level, we adjust for IGC, which has been demonstrated to inflate type-I error if mistakenly ignored. Using gene-level residuals to estimate IGC, we produce set-level *p*-values that preserve type-I error and improve power over alternative approaches.

The ability of TWO-SIGMA-G to include random effect terms at the gene-level provides a distinguishing factor from many methods for gene set analysis. Such random effects can improve inference at the gene-level substantially for some genes [38]. However, if only interested in set-level inference, our simulations suggest that statistical inference remains valid when excluding gene-level random effects and reducing computational burden as a result. When gene-level inference is also of interest, it is likely desirable to fall back on including random effect terms into the regression modelling framework. However, we suggest that inference in real data analyses is likely not influenced greatly at the set-level by the presence or absence of gene-level random effects (Supplementary Figure S29).

TWO-SIGMA-G is implemented in the `twosigma` R package (https://github.com/edvanburen/twosigma), which is computationally efficient and allows for parallelization. To benchmark computational performance, we ran a modified version of our HIV data analysis, testing for a treatment effect of HIV pooled over all cell types. This modification to a one degree of freedom hypothesis allows us to test identical hypotheses in TWO-SIGMA-G, MAST, and CAMERA to provide a fairer comparison of computation. Using three computing cores on a MacBook Pro laptop, the methods had the following respective runtimes: 33.2 minutes for TWO-SIGMA-G, 33.5 minutes for MAST (25 bootstrap replications), and 5 seconds for CAMERA. TWO-SIGMA-G shows slightly improved yet nearly identical computational performance to MAST in the presence of the other advantages for performing gene set testing in scRNA-seq data described throughout this paper.

Unlike bulk RNA-seq data, many genes are often uncaptured or fail to survive filtering in scRNA-seq data. In gene set analysis, we must therefore assume that a gene set can be represented by the genes that exist in the dataset. In both the HIV and Alzheimer’s datasets, we typically have around 40% representation regardless of set size after gene filtering (Supplementary Figure S30). Given the biologically meaningful and interpretable results we presented, the absence of these genes does not seem to threaten the ability of scRNA-seq gene set analyses to contribute new biological insights.

## 5 Data Availability

Both datasets analyzed in this manuscript are publicly available. The HIV dataset is available at the Gene Expression Omnibus under accession GSE148796. The Alzheimer’s dataset is available upon completion of a data usage agreement at the Rush Alzheimer’s Disease Center (RADC) Research Resource Sharing Hub (https://www.radc.rush.edu/docs/omics.htm) under “snRNA-seq PFC.” TWO-SIGMA-G is implemented in the function `twosigmag` in the `twosigma R` package, which is freely available on GitHub at https://github.com/edvanburen/twosigma.

## 7 Acknowledgements

None at this time.

