## supplement for "TWO-SIGMA-G: A New Competitive Gene Set Testing Framework for scRNA-seq Data Accounting for Inter-Gene and Cell-Cell Correlation"

### S1 Gene Set Simulation Procedure

#### S1.1 Zero-Inflated Negative Binomial Distribution

We use the zero-inflated negative binomial (ZINB) distribution to simulate gene-level expression data. For a given gene, let $i$ index the samples sequenced and $j$ index the $n_i$ single cells from sample $i$. Consider the following parameterization of the negative binomial probability mass function (p.m.f.) at a non-negative integer $y_{ij}$ corresponding to the observed read count:

$$\begin{aligned}\Pr(Y_{ij} = y_{ij}) &= f(y_{ij}; \mu_{ij}, \phi) \\ &= \frac{\Gamma\Big(y_{ij} + \phi\Big)}{\Gamma(y_{ij}+1)\Gamma\Big(\phi\Big)} \left(\frac{1}{1+\frac{1}{\phi}\mu_{ij}}\right)^{\phi} \left(\frac{\frac{1}{\phi}\mu_{ij}}{1+\frac{1}{\phi}\mu_{ij}}\right)^{y_{ij}} \\ &,\ y_{ij} = 0, 1, 2, \ldots\end{aligned}$$

With this parameterization, $\mathrm{E}(Y_{ij}) = \mu_{ij}$ and $\mathrm{Var}(Y_{ij}) = \mu_{ij} + \frac{1}{\phi}\mu_{ij}^2$, such that $\phi$ is the overdispersion parameter ($\phi > 0$). Let $p_{ij}$ and $\mu_{ij}$ be the probability of drop-out and the mean read count conditional on not being dropped-out for cell $j$ from sample $i$, respectively. The p.m.f. of one observation $Y_{ij}$ under the ZINB($\mu_{ij}$, $p_{ij}$,$\phi$) distribution is given by:

$$\begin{aligned}\mathrm{P}(Y_{ij} = 0) &= p_{ij} + (1 - p_{ij}) f(0; \mu_{ij}, \phi) \\ \mathrm{P}(Y_{ij} = y_{ij}) &= (1 - p_{ij}) f(y_{ij}; \mu_{ij}, \phi), y_{ij} = 1, 2, 3, \ldots\end{aligned}$$

We assume the parameters $p_{ij}$ and $\mu_{ij}$ come from the following mixed effects regression model:

$$\begin{aligned}\mathrm{logit}(p_{ij}) &= \boldsymbol{z_{ij}^T}\boldsymbol{\alpha} + a_i, a_i \sim N(0, \sigma_a^2) \\ \log(\mu_{ij}) &= \boldsymbol{x_{ij}^T}\boldsymbol{\beta} + b_i, b_i \sim N(0, \sigma_b^2),\ \text{assume } a_i \perp\!\!\!\perp b_i \end{aligned} \quad (1)$$

Here, $\boldsymbol{\alpha}$ and $\boldsymbol{\beta}$ are fixed effect coefficient vectors with corresponding vectors of covariates $\boldsymbol{z_{ij}}$ and $\boldsymbol{x_{ij}}$. $a_i$ and $b_i$ are sample-specific random intercept terms following normal distributions with mean 0 and variances $\sigma_a^2$ and $\sigma_b^2$, respectively.

#### S1.2 Procedure to Simulate Correlated Gene Sets

To simulate correlated gene sets, with varying magnitudes of correlation designed to represent the diversity seen in real data, we employ the following simulation procedure:

1. Simulate independent genes

   (a) Simulate population of 10,000 cells from 100 samples using one simulated design matrix with covariates simulated as follows: (i) a binary treatment effect of interest ("trt", $\sim$ Binomial(0.4)) and (ii) additional covariates corresponding to a continuous confounder (e.g., "age", $\sim$ Unif(20,60)), and a cell-level measure of the CDR ($\sim$ Beta(3,6)). These distributions for the design matrix will be re-used for generating more design matrices in the steps below.

   (b) Simulate values for $\boldsymbol{\alpha}$, $\boldsymbol{\beta}$ (from equation (1)) for covariates from (a) using the following model:

$$\begin{aligned} \boldsymbol{z}_{\boldsymbol{ij}}^T\boldsymbol{\alpha} &= \alpha_0 + \alpha_1\text{trt}_i + \alpha_2\text{age}_i + \alpha_3\text{CDR}_j \\ \boldsymbol{x}_{\boldsymbol{ij}}^T\boldsymbol{\beta} &= \beta_0 + \beta_1\text{trt}_i + \beta_2\text{age}_i + \beta_3\text{CDR}_j \end{aligned} \quad (2)$$

   - Intercepts $\alpha_0 = 1$ and $\beta_0 = 2$ fixed to ensure drop-out percentages and data scale comparable
   - Values $\alpha_1$ and $\beta_1$ for binary treatment effect are equal to zero (null hypothesis) or non-zero (alternative), with the following relations to presented results:

| ($\alpha_1$, $\beta_1$) | Figures | Comments |
|---|---|---|
| (0,0) | 2, S1, S2, S3 | Null Scenarios |
| (0.03, 0.03) | 4, S4, S7 | Comparisons to GSEA, MAST & CAMERA |
| 50% = (0.03, 0.03), 50% = (0.06, 0.06) | S8 | "Mixed DE". Comparisons to GSEA, MAST & CAMERA |
| (0.03, 0.20) | 3, S5, S6, S9, S10 | Comparisons to iDEA, fGSEA, & PAGE |

   - Values $\alpha_2$, $\alpha_3$, $\beta_2$, and $\beta_3$ either (i) equal to zero (simplifies the gene-gene correlation structure when introducing IGC in Step 2) or (ii) simulated from Uniform(-1.5,1.5) (See Supplementary Table S1)

   (c) Set $\sigma_a^2 = \sigma_b^2 = 0$ to create no gene-level RE terms or $\sigma_a^2 = \sigma_b^2 = 0.1$ to create gene-level RE terms

   (d) Using (a), (b), and (c) calculate parameters $p_{ij}$ and $\mu_{ij}$ using equation 1

   (e) Simulate $Y_{ij}$ from the ZINB distribution with parameters from (d) and setting $\phi$ = 0.1.

   (f) Simulate 1,000 independent genes without gene-level random effect terms and 300 independent genes with gene-level random effect terms for two conditions, i.e., when there are additional covariates and when there are no additional covariates (Supplementary Table S1).

Supplementary Table S1: Shows the six different settings used to simulate data for gene set simulations. $w_2$ controls how two genes are correlated, with a larger $w_2$ corresponding to lower IGC. "A.C." refers to the presence of additional covariates, age and CDR, in addition to the treatment effect.

| $w_2$ | A.C. | $(\alpha_2, \alpha_3, \beta_2, \beta_3)$ |
|---|---|---|
| 3 | No | $\equiv 0$ |
| 5 | No | $\equiv 0$ |
| 10 | No | $\equiv 0$ |
| 3 | Yes | $\sim$ Unif$(-1.5, 1.5)$ |
| 5 | Yes | $\sim$ Unif$(-1.5, 1.5)$ |
| 10 | Yes | $\sim$ Unif$(-1.5, 1.5)$ |

2. Generate correlated gene sets of size 30 (Set sizes of 30 were chosen to approximately correspond to the size of a "typical" set in the MsigDB c2 collection).

   (a) For each independent gene, $Y_{indep}$, add noise from NB distribution using pre-specified, fixed parameters, $\mu_{perm}$, $\phi_{perm}$, and $w_2$ to create 29 correlated genes $Y_{corr,j}$, for $j = 1, ..., 29$:

   $$Y_{corr,j} = \text{round}(0.5 * Y_{indep} + w_2 * NB(3, 2.5))$$

   - Weight NB noise by $w_2$ to control the amount of correlation (larger $w_2$ means lower IGC) and change mean patterns across scenarios (See Supplementary Table S1 below for more details)
   - If gene is under the alternative, add additional noise $w_3 * NB(3, 2.5)$ to preserve signal and correlation structure ($w_3$ taken as 0.15 in "mixed DE" alternatives and 0.1 otherwise)

   (b) Randomly set some non-zero counts to zero to keep the proportion of zeros the same in correlated and independent gene

   - Ensures that proportion of zeros alone does not drive significant results

3. Vary magnitude of IGC in 2(a) by taking $w_2$ from various values in Supplementary Table S1. Each of the six settings is simulated both with (300 times to create 300 gene sets) and without gene-level random effect terms (1,000 times to create 1,000 gene sets). When we test one set of 30 correlated genes in TWO-SIGMA-G, the other genes (29,999 genes when there are no random effects, 8,999 genes when there are random effects) are used as candidate genes for reference sets in the gene set test.

4. Repeat the above steps 1-3 for 10 times. For each time, a different random seed is used to create a new design matrix, allowing for more randomness of the covariates in design matrix and thus more randomness in the the gene expression data. In total, type-I error simulations were repeated 3,000 times without RE and 900 times with RE, and power simulations were repeated 600 times without RE and 300 times with RE.

### S2 Additional Type-I Error and Power Results

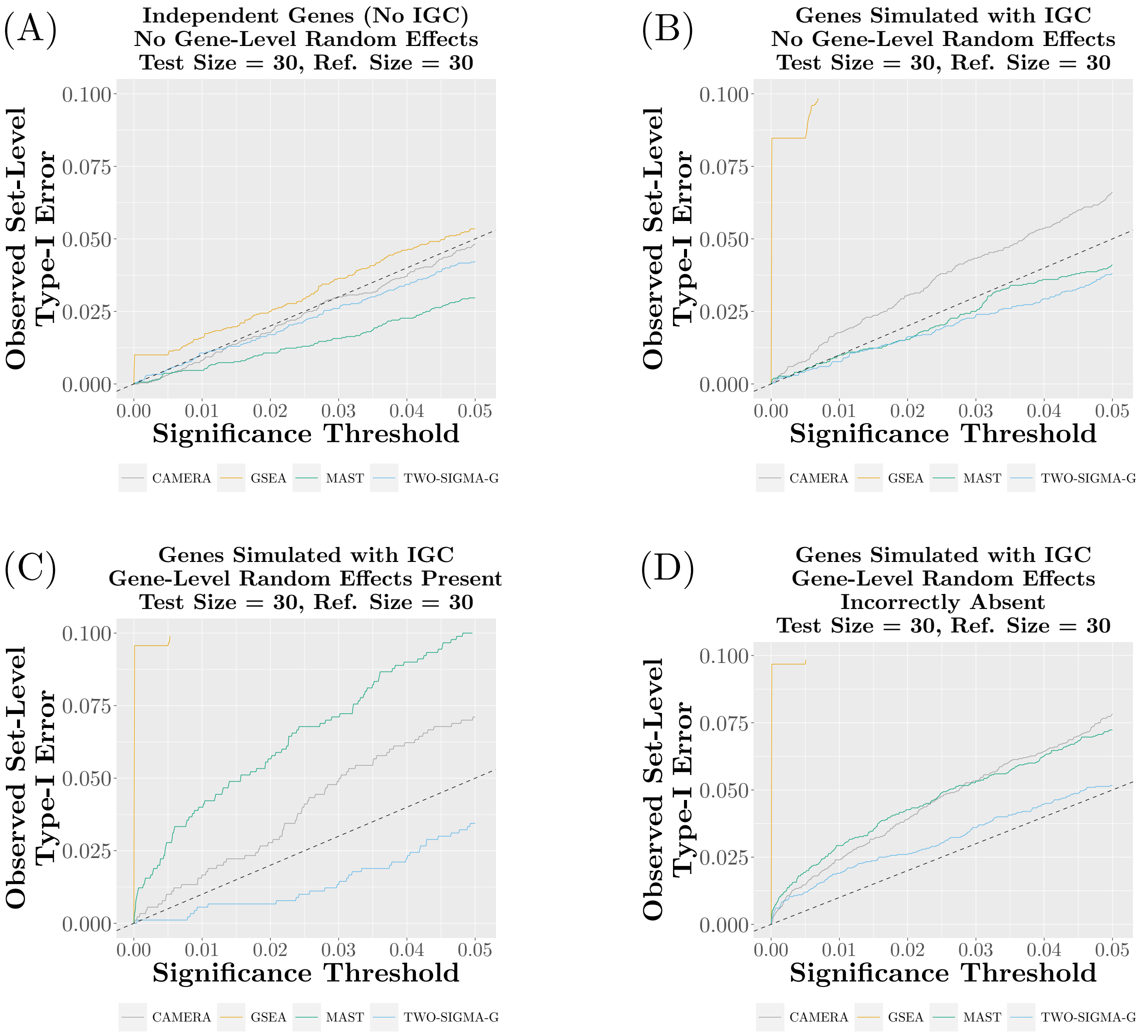


Supplementary Figure S1: **Type-I error performance of CAMERA, GSEA, MAST, and TWO-SIGMA-G as significance threshold varies from 0 to .05**. Each panel varies the existence of IGC between genes in the test set and the presence of gene-level random effect terms in gene-level model (CAMERA and GSEA never include gene-level random effect terms). Each plot combines six different settings, and 10 replicates per setting, which vary both the magnitude of the average inter-gene correlation (where applicable) in the test set and the nature of the correlation structure via the introduction of other individual-level covariates. See the Methods section of the main text for more details regarding the simulation procedure.

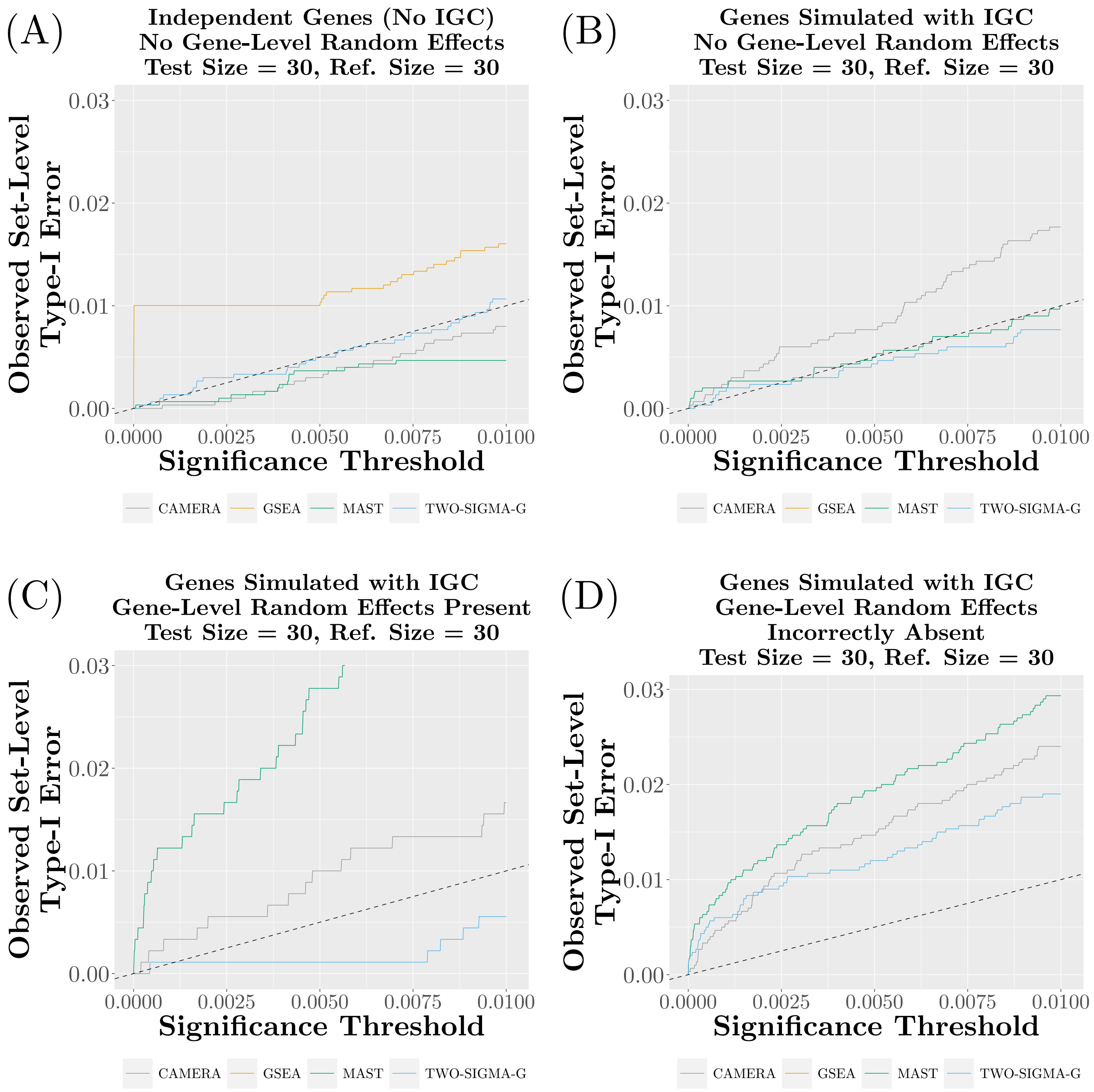


Supplementary Figure S2: **Type-I error performance of CAMERA, MAST, and TWO-SIGMA-G as significance threshold varies from 0 to .01**. Each panel varies the existence of IGC between genes in the test set and the presence of gene-level random effect terms in gene-level model (CAMERA and GSEA never include gene-level random effect terms). Each plot combines six different settings, and 10 replicates per setting, which vary both the magnitude of the average inter-gene correlation (where applicable) in the test set and the nature of the correlation structure via the introduction of other individual-level covariates. See the Methods section of the main text for more details regarding the simulation procedure.

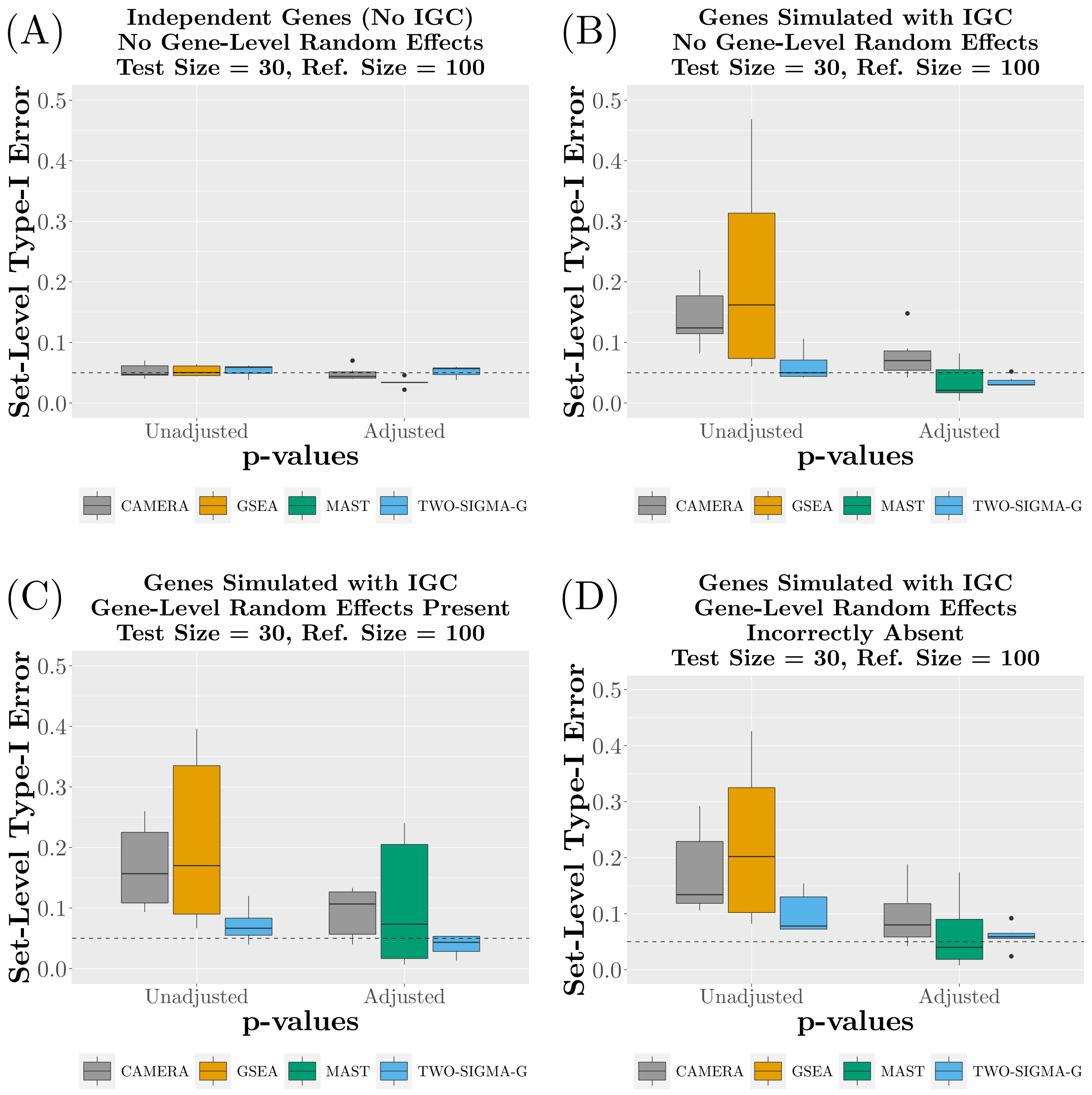


Supplementary Figure S3: **Type-I error performance of CAMERA, GSEA, MAST, and TWO-SIGMA-G using a reference set size of 100 genes**. Each panel varies the existence of IGC between genes in the test set and the presence of gene-level random effect terms in gene-level model (CAMERA and GSEA never include gene-level random effect terms). Within each panel, both unadjusted and adjusted set-level $p$-values are plotted (unadjusted $p$-values are unavailable for MAST). See Supplementary Section S1 and the Methods section of the main text for more details regarding the simulation procedure.

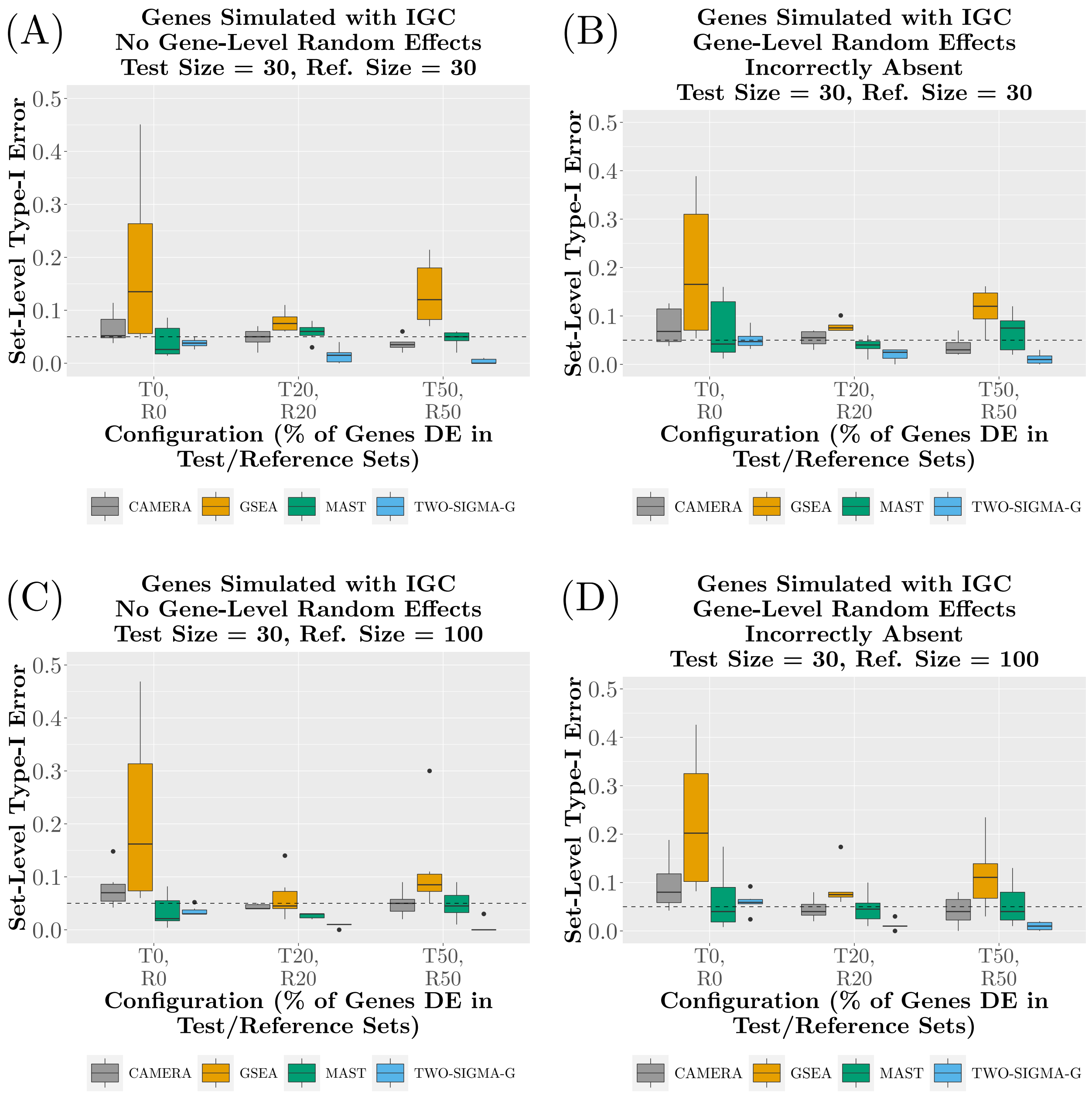


Supplementary Figure S4: **Type-I error performance of TWO-SIGMA-G, CAMERA, GSEA, and MAST for various set-level null hypotheses**. Each panel varies the reference set size and the presence of gene-level random effect terms in gene-level model. Scenarios along the $x$-axis of each panel vary the percentage of genes that are differentially expressed (with the same effect size) in the test and reference sets. See Supplementary Section S1 and the Methods section of the main text for more details regarding the simulation procedure.

(A)

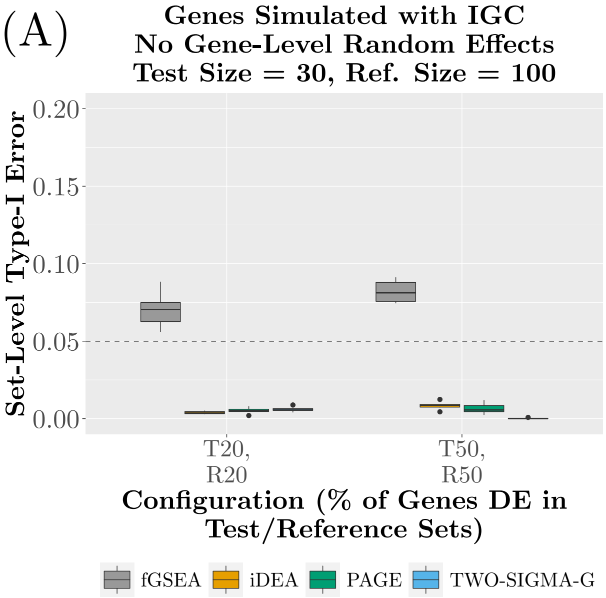


Supplementary Figure S5: **Type-I error performance of iDEA, PAGE, fGSEA, and TWO-SIGMA for various set-level null hypotheses for genes simulated with IGC**. Reference set sizes of 100 are shown. See Supplementary Section S1 and the Methods section of the main text for more details regarding the simulation procedure.

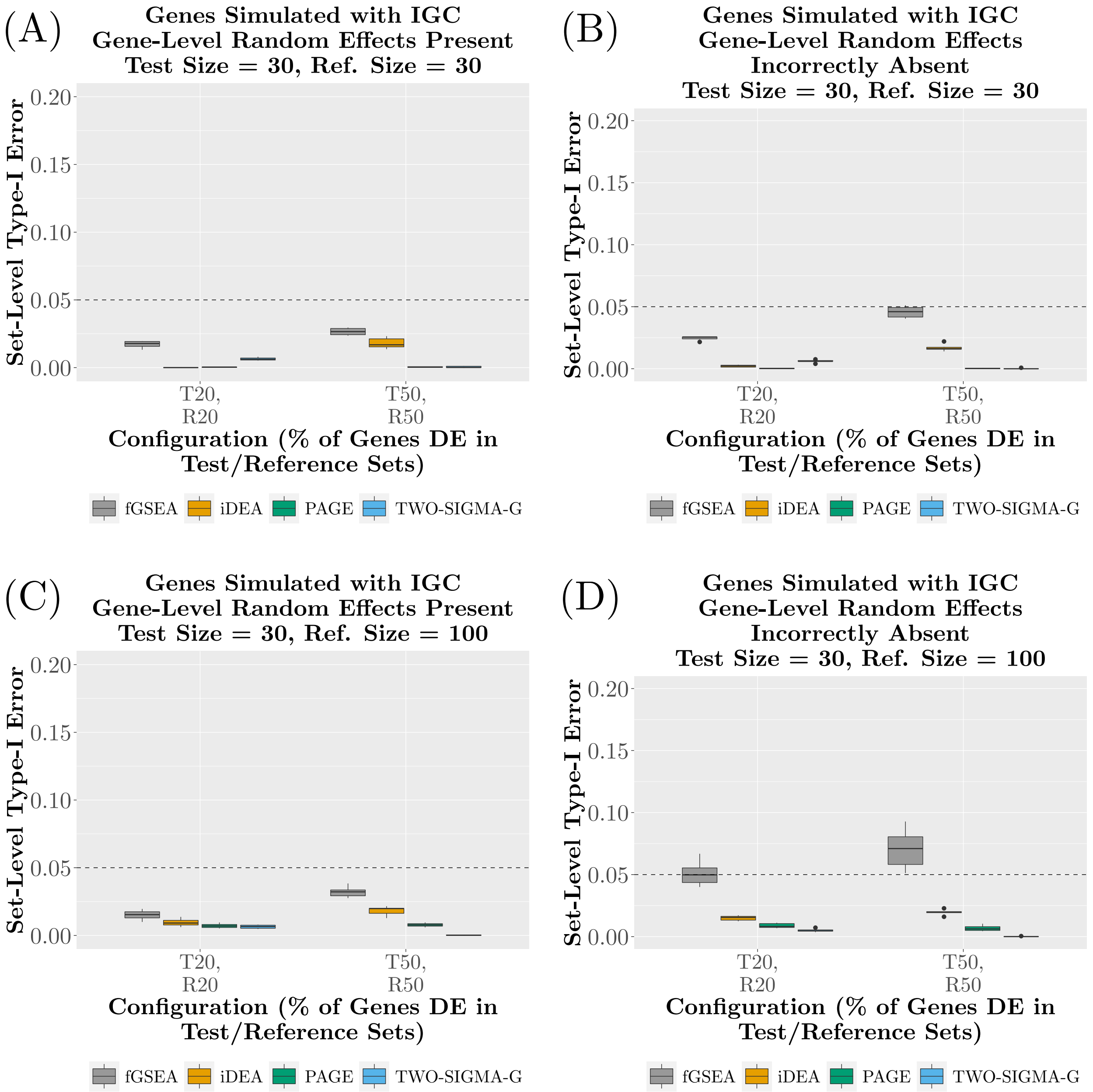


Supplementary Figure S6: **Type-I error performance of iDEA, PAGE, fGSEA, and TWO-SIGMA for various set-level null hypotheses with gene-level random effect terms**. See Supplementary Section S1 and the Methods section of the main text for more details regarding the simulation procedure.

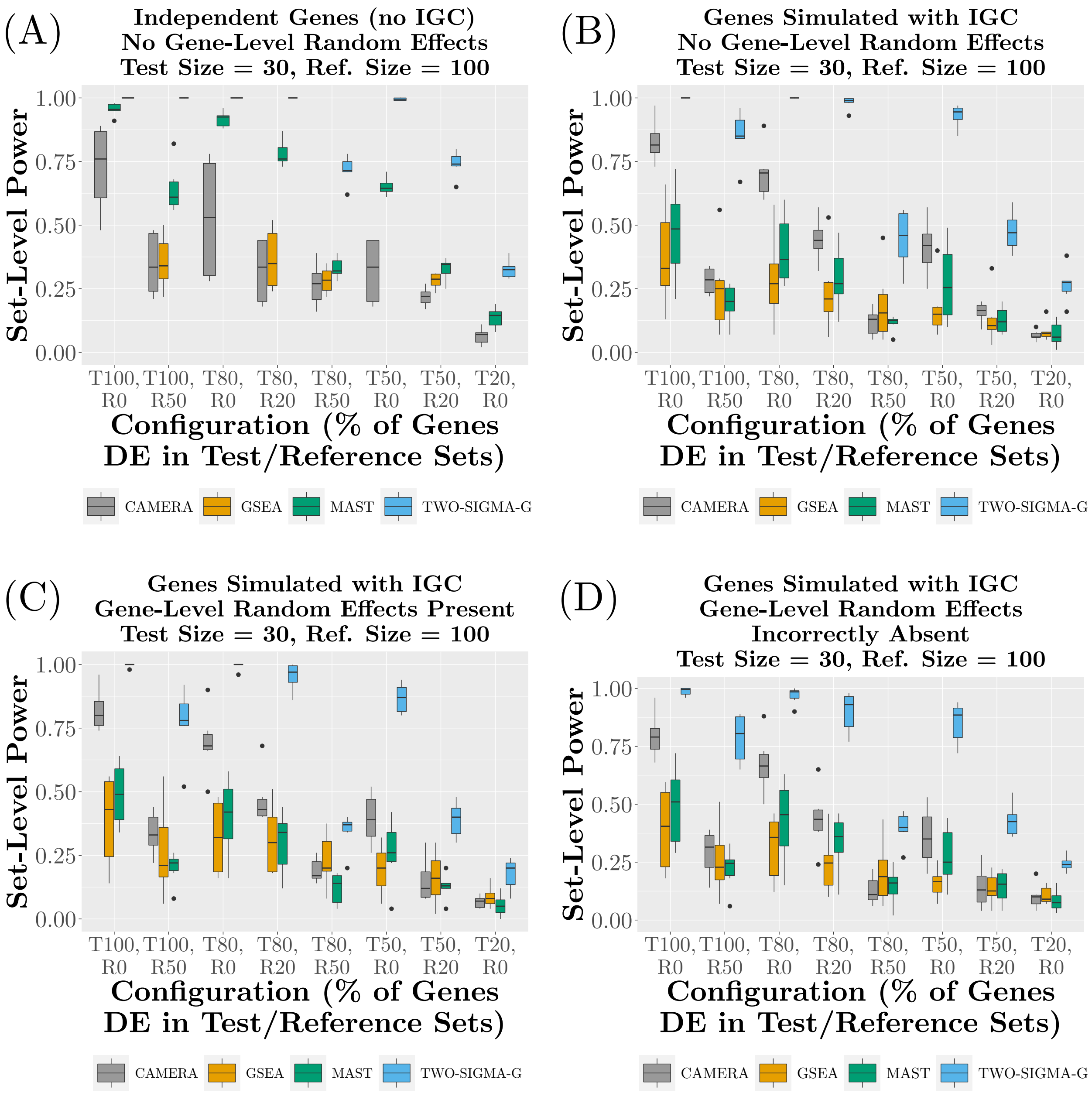


Supplementary Figure S7: **Set-level power of CAMERA, GSEA, MAST, and TWO-SIGMA-G using a reference set size of 100 genes (corresponds to Figure 3 in main text)**. Each panel varies the existence of IGC between genes in the test set and the presence of gene-level random effect terms in gene-level model (CAMERA and GSEA never include gene-level random effect terms). Scenarios along the $x$-axis of each panel vary the percentage of genes that are differentially expressed (with the same effect size) in the test and reference sets. For example, “T80,R50” corresponds to the configuration under the alternative hypothesis in which 80% of test set genes are DE and 50% of reference set genes are DE. See Supplementary Section S1 and the Methods section of the main text for more details regarding the simulation procedure.

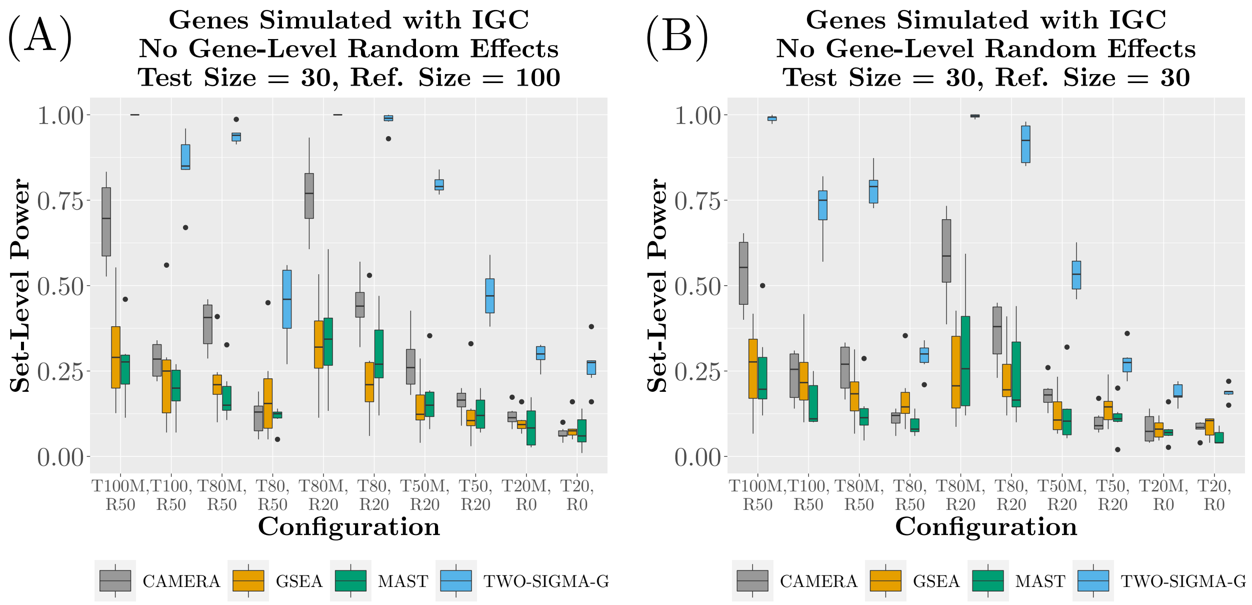


Supplementary Figure S8: **Set-level power of CAMERA, GSEA, MAST, and TWO-SIGMA-G using differing DE magnitudes at the gene-level**. Genes are simulated with IGC, and reference set sizes of 100 and 30 are used for gene set testing. Scenarios along the $x$-axis of each panel vary the percentage of genes that are differentially expressed (with the same effect size) in the test and reference sets. For example, "T80,R50" corresponds to the configuration under the alternative hypothesis in which 80% of test set genes are DE and 50% of reference set genes are DE. Within some test sets, the amount of DE is mixed: with 50% of genes having twice as large of an effect size as the other half (see, e.g., "T100M, R50"). See Supplementary Section S1 and the Methods section of the main text for more details regarding the simulation procedure.

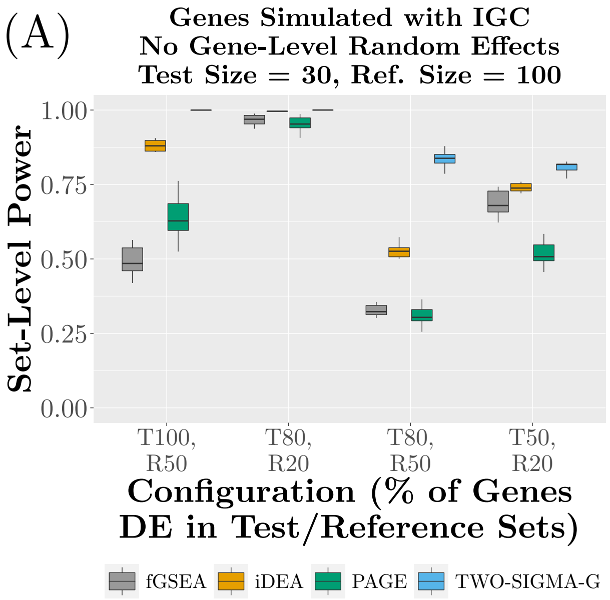


Supplementary Figure S9: **Set-level power of iDEA, PAGE, fGSEA, and TWO-SIGMA-G at the gene-level**. Genes are simulated with IGC using reference set sizes of 100. Scenarios along the $x$-axis of each panel vary the percentage of genes that are differentially expressed (with the same effect size) in the test and reference sets. For example, "T80,R50" corresponds to the configuration under the alternative hypothesis in which 80% of test set genes are DE and 50% of reference set genes are DE. Because iDEA performed poorly in scenarios involving "R0", they were excluded. See Supplementary Section S1 and the Methods section of the main text for more details regarding the simulation procedure.

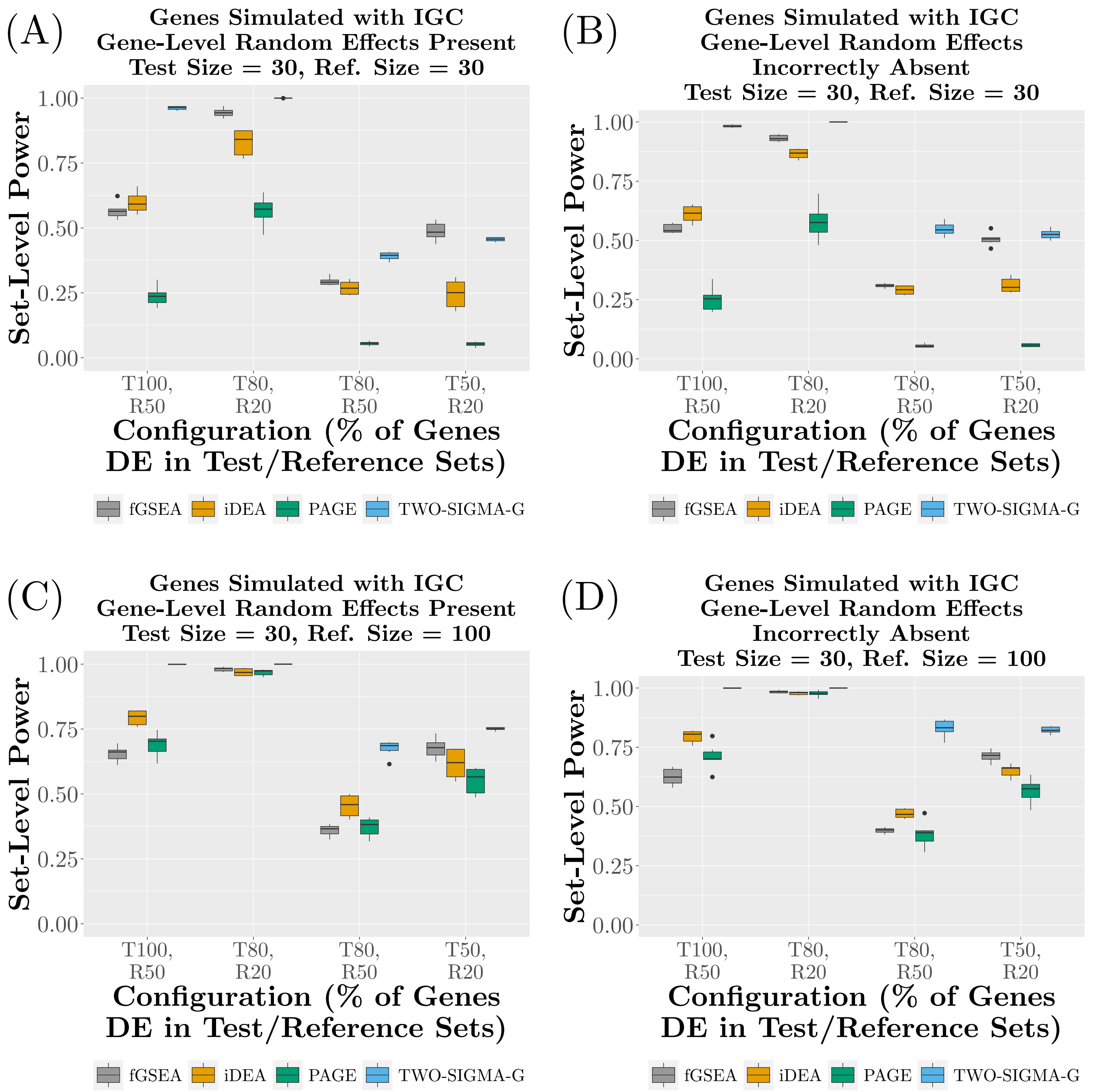


Supplementary Figure S10: **Set-level power of iDEA, PAGE, fGSEA, and TWO-SIGMA-G when gene-level random effects are included**. Scenarios along the $x$-axis of each panel vary the percentage of genes that are differentially expressed (with the same effect size) in the test and reference sets. For example, "T80,R50" corresponds to the configuration under the alternative hypothesis in which 80% of test set genes are DE and 50% of reference set genes are DE. Because iDEA performed poorly in scenarios involving "R0", they were excluded. See Supplementary Section S1 and the Methods section of the main text for more details regarding the simulation procedure.

### S3 Additional HIV Data Results

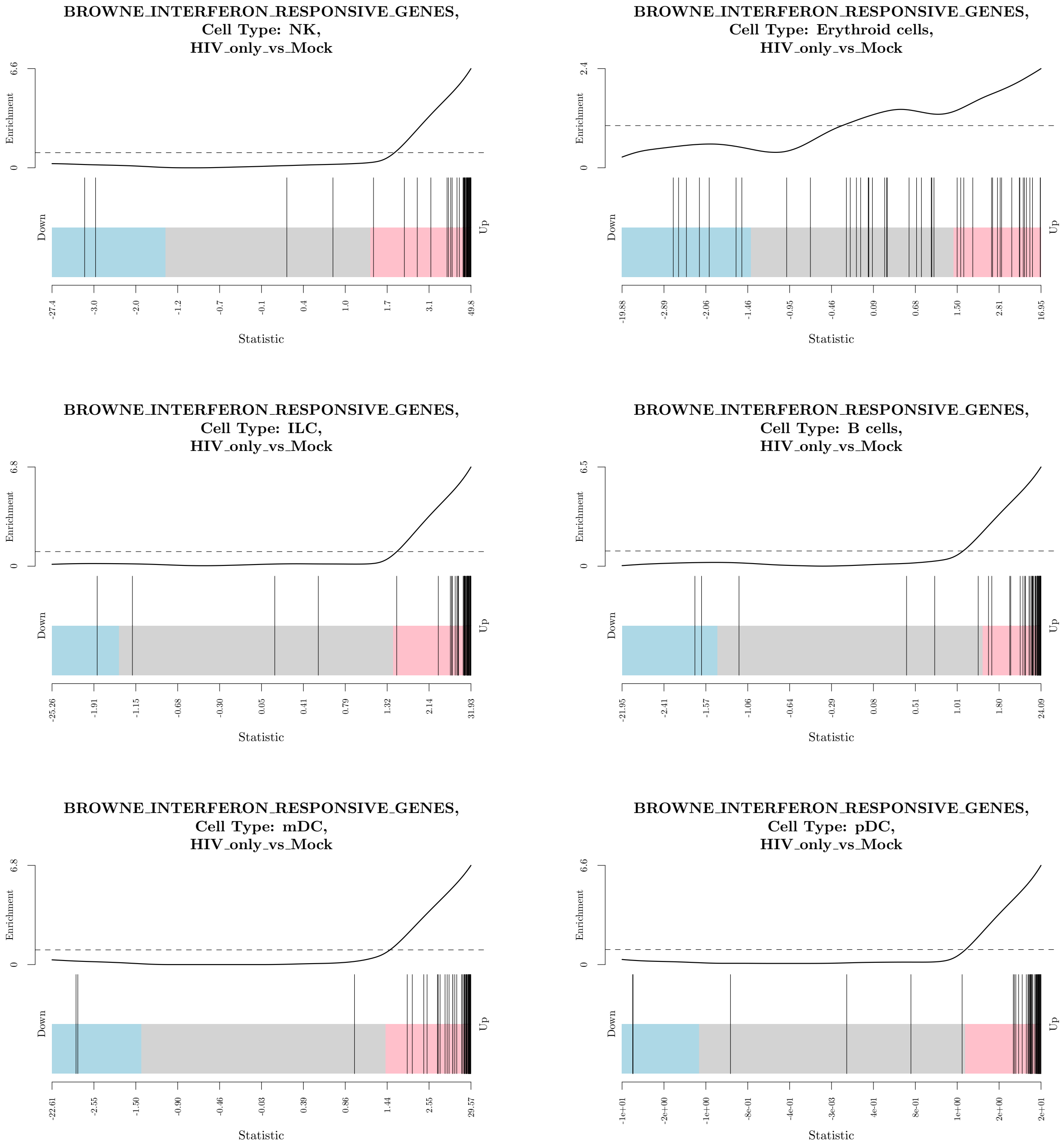


Supplementary Figure S11: **Barcode plots showing gene-level statistics from TWO-SIGMA-G for the six most common cell types in the HIV dataset for the gene set "BROWNE_INTERFERON_RESPONSIVE_GENES."** The x-axis shows the concentration of gene-level statistics (each bar represents one gene-level statistic), and the enrichment shown in the y-axis demonstrates that gene-level statistics tend to be overwhelmingly positive and large in magnitude for all cell types except Erythroid cells.

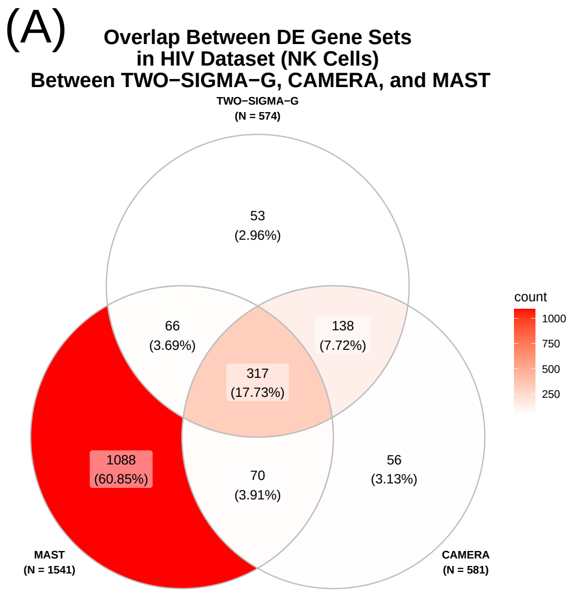


Supplementary Figure S12: **Venn diagram showing set-level overlap across methods among all DE gene sets before FDR adjustment for NK cells in the HIV data.**

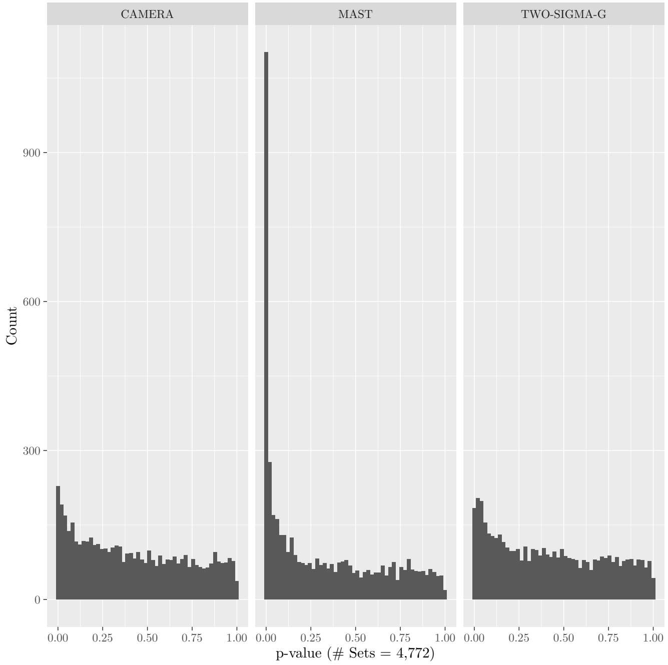


Supplementary Figure S13: **Histogram showing set-level *p*-values from NK cells for MAST, TWO-SIGMA-G, and CAMERA in the HIV data.**

### S4 Additional Alzheimer's Results: Comparing Early Stage AD to Control

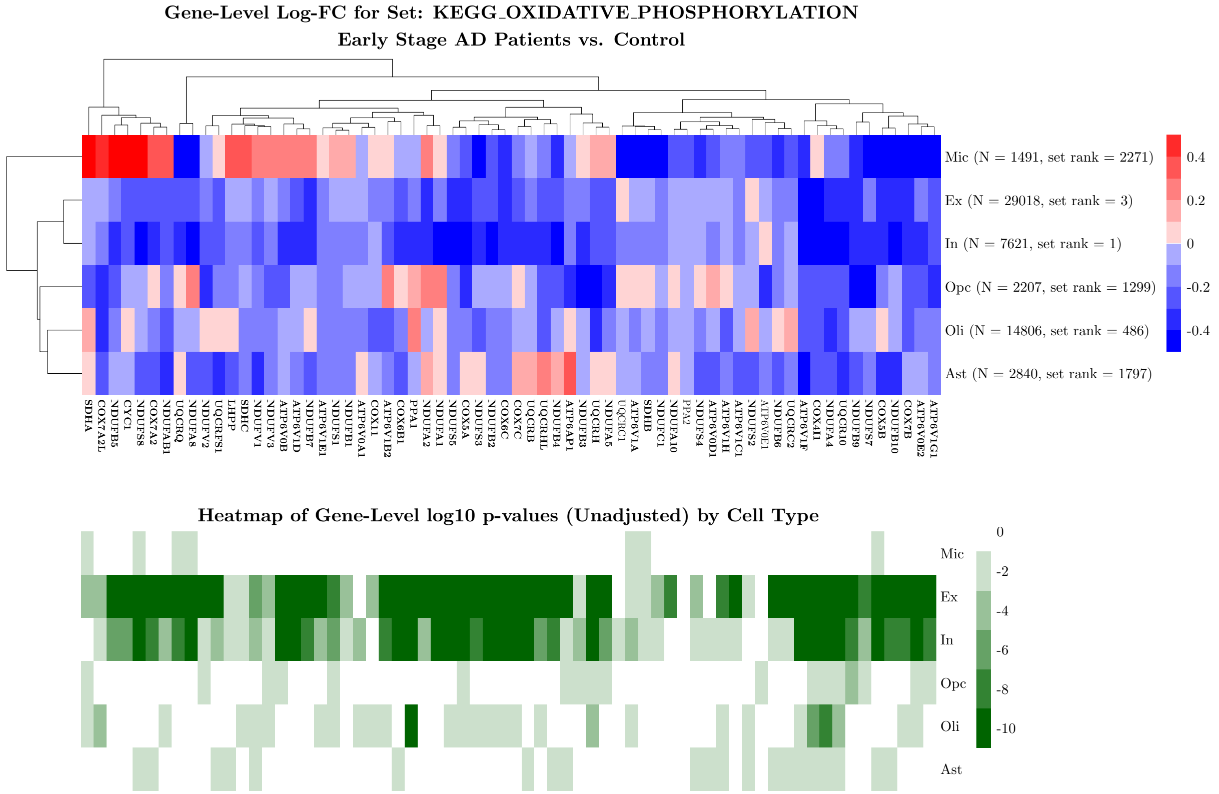


Supplementary Figure S14: **Variation in gene-level significance at the cell type level for genes in the KEGG_OXIDATIVE_PHOSPHORYLATION pathway comparing early stage AD patients to controls**. Gene names that are bolded are signifcant over all cell types after FDR-adjustment of the Fisher's method $p$-value.

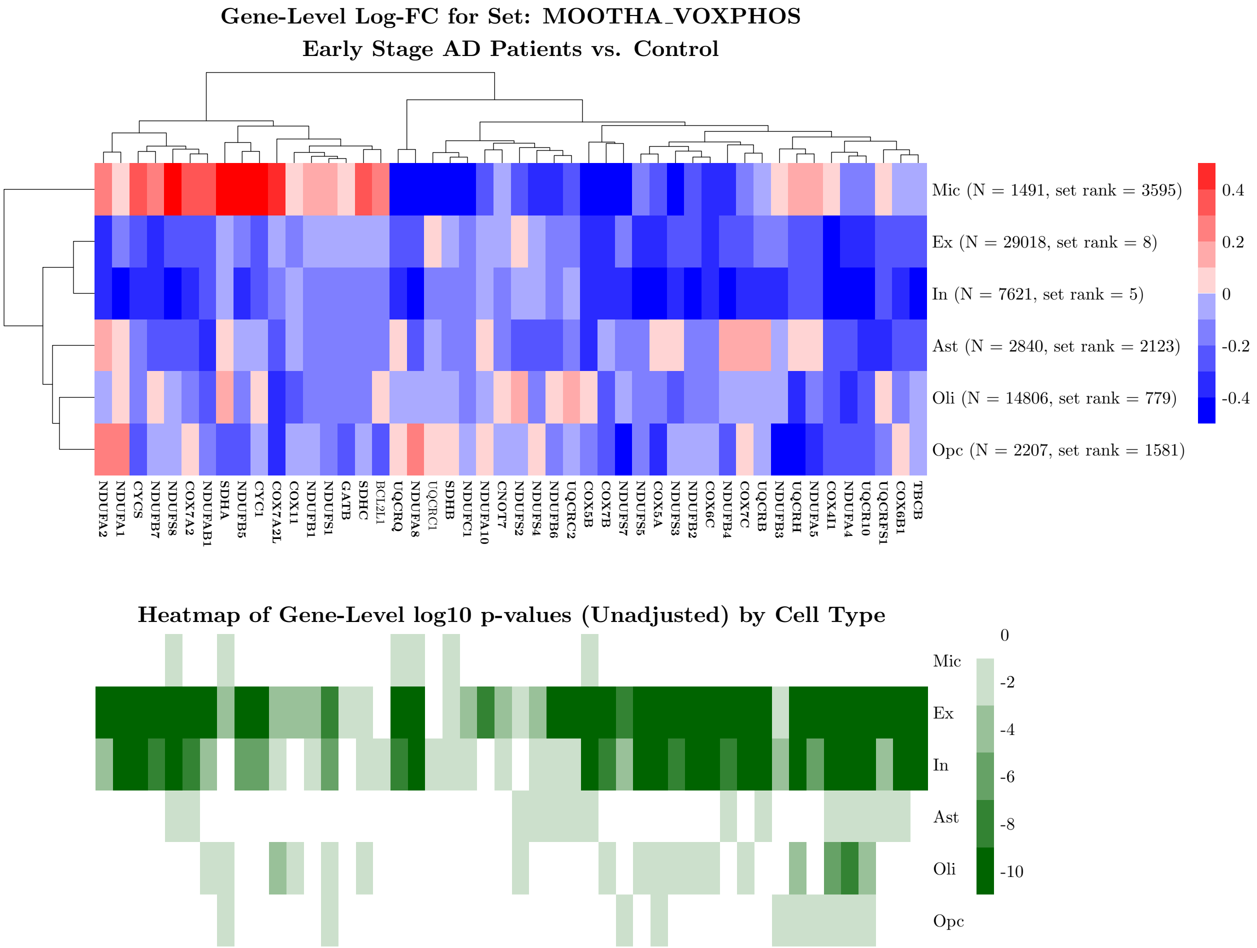


Supplementary Figure S15: **Variation in gene-level significance at the cell type level for genes in the MOOTHA_VOXPHOS pathway comparing early stage AD patients to controls**. Gene names that are bolded are signifcant over all cell types after FDR-adjustment of the Fisher's method $p$-value.

### S5 Additional Alzheimer's Results: Comparing Late to Early Stage AD

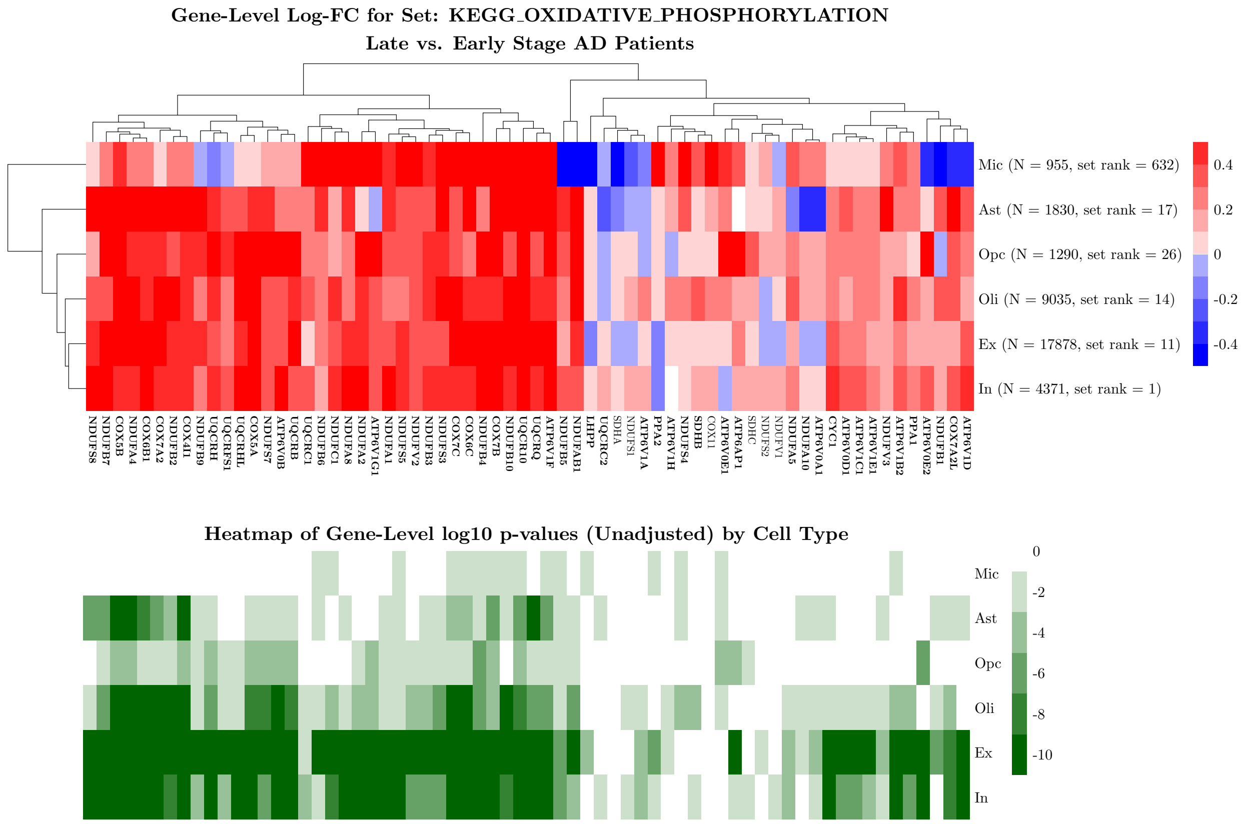


Supplementary Figure S16: **Variation in gene-level significance at the cell type level for genes in the KEGG_OXIDATIVE_PHOSPHORYLATION pathway comparing late stage AD patients to early stage AD patients**. Gene names that are bolded are signifcant over all cell types after FDR-adjustment of the Fisher's method $p$-value.

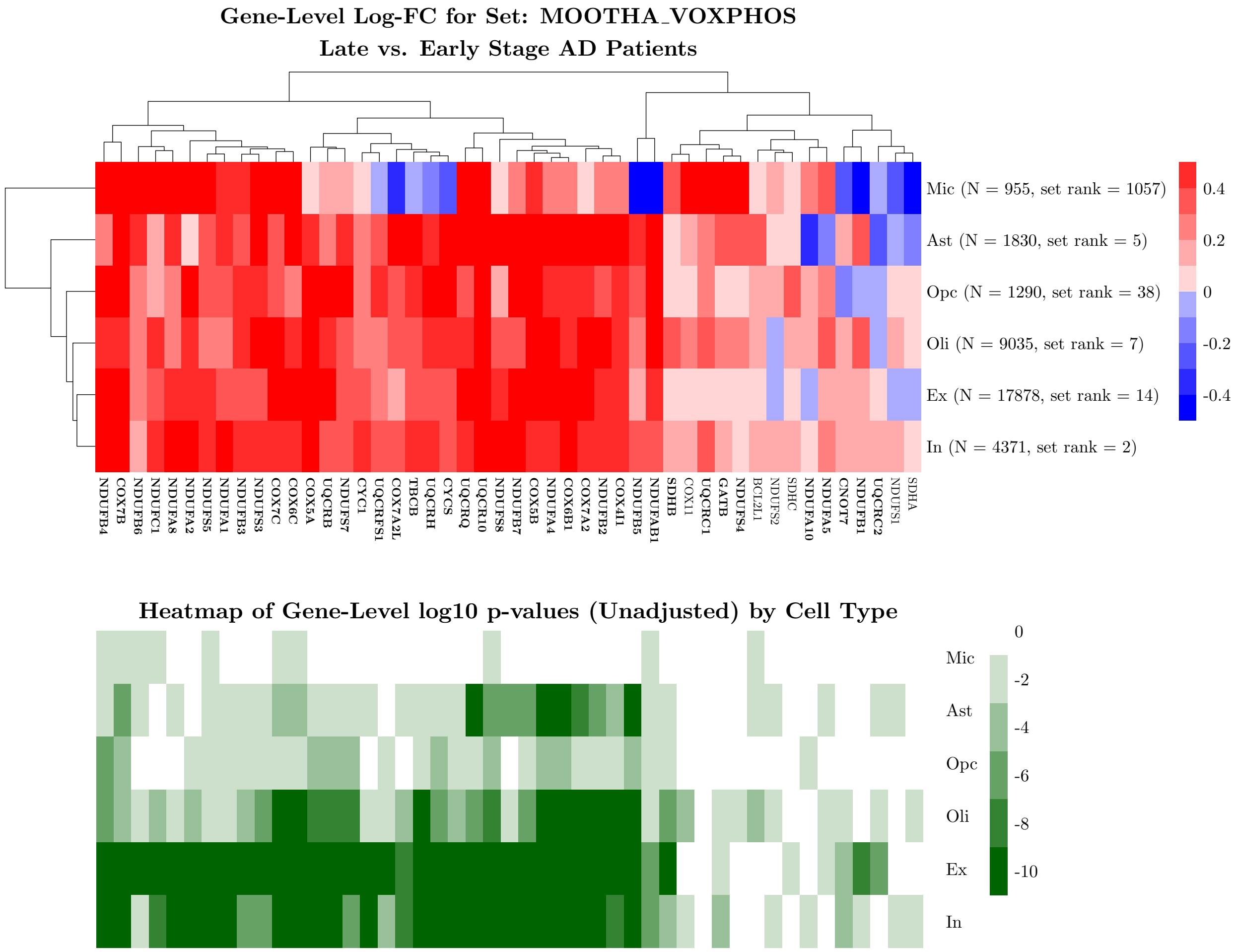


Supplementary Figure S17: **Variation in gene-level significance at the cell type level for genes in the MOOTHA_VOXPHOS pathway comparing late stage AD patients to early stage AD patients**. Gene names that are bolded are signifcant over all cell types after FDR-adjustment of the Fisher's method $p$-value.

### S6 Additional Alzheimer's Results: Comparing Late Stage AD to Controls

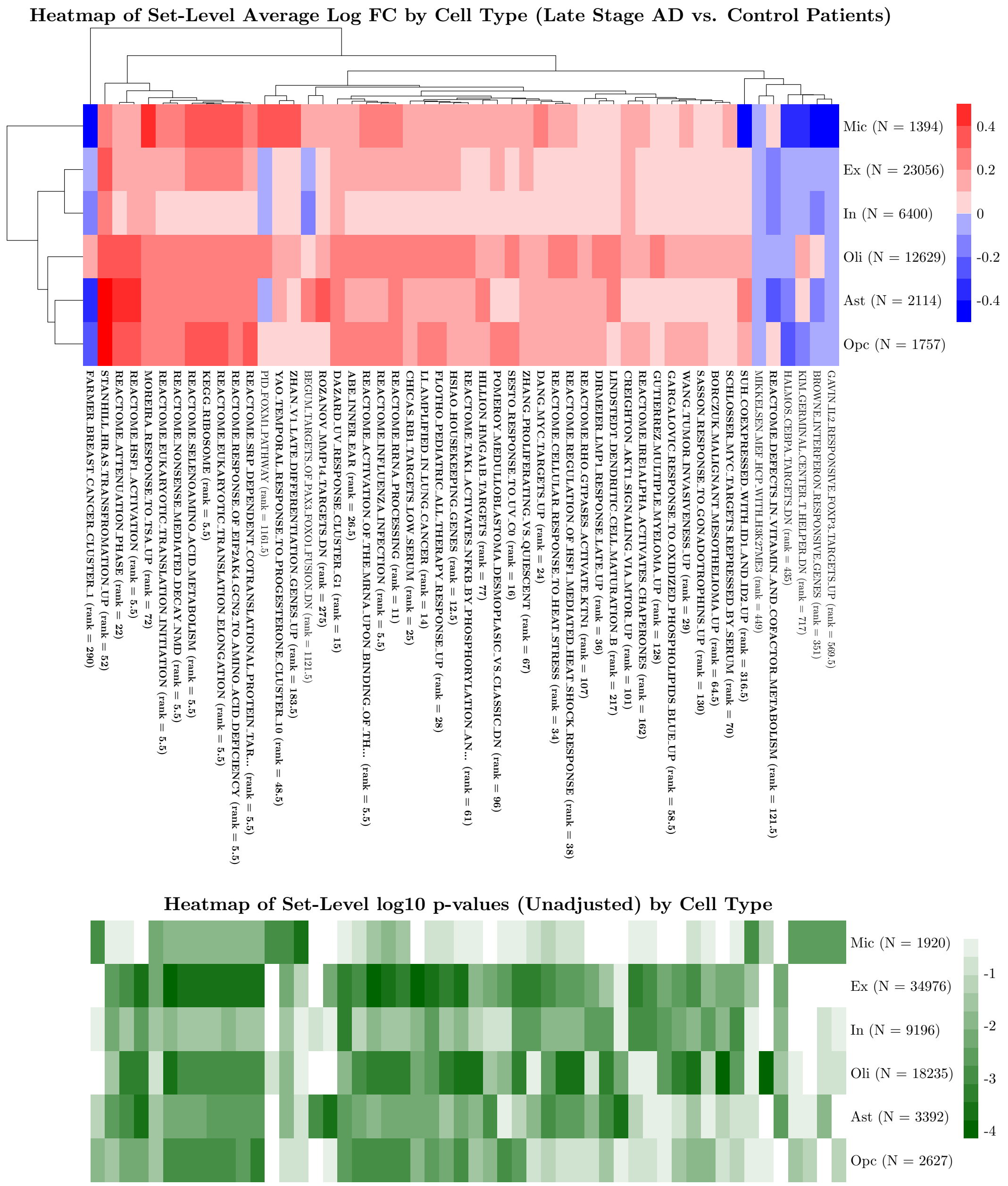


Supplementary Figure S18: **Heatmap of the most significant gene sets (and their corresponding p-values) comparing late state AD patients to controls by cell type**. Sets plotted are among the top 10 in significance for at least once cell type. Sets in bold are significant over all cell types after FDR-adjustment of the Fisher's method *p*-value.

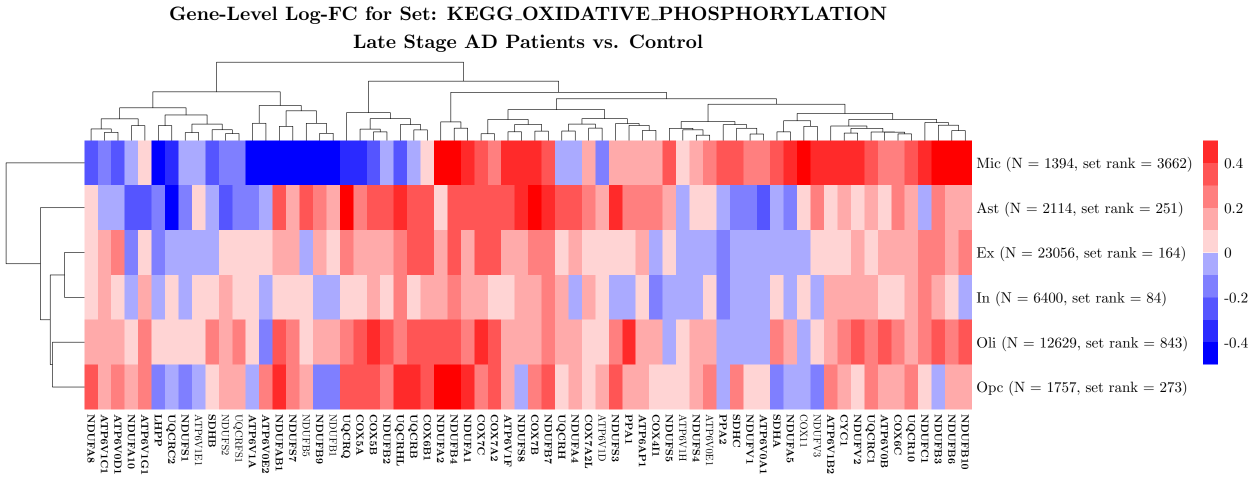


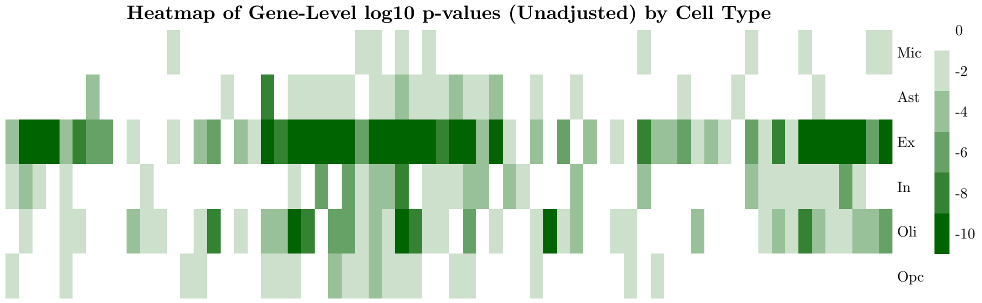


Supplementary Figure S19: **Variation in gene-level significance at the cell type level for genes in the KEGG_OXIDATIVE_PHOSPHORYLATION pathway comparing late stage AD patients to controls**. Gene names that are bolded are signifcant over all cell types after FDR-adjustment of the Fisher's method $p$-value.

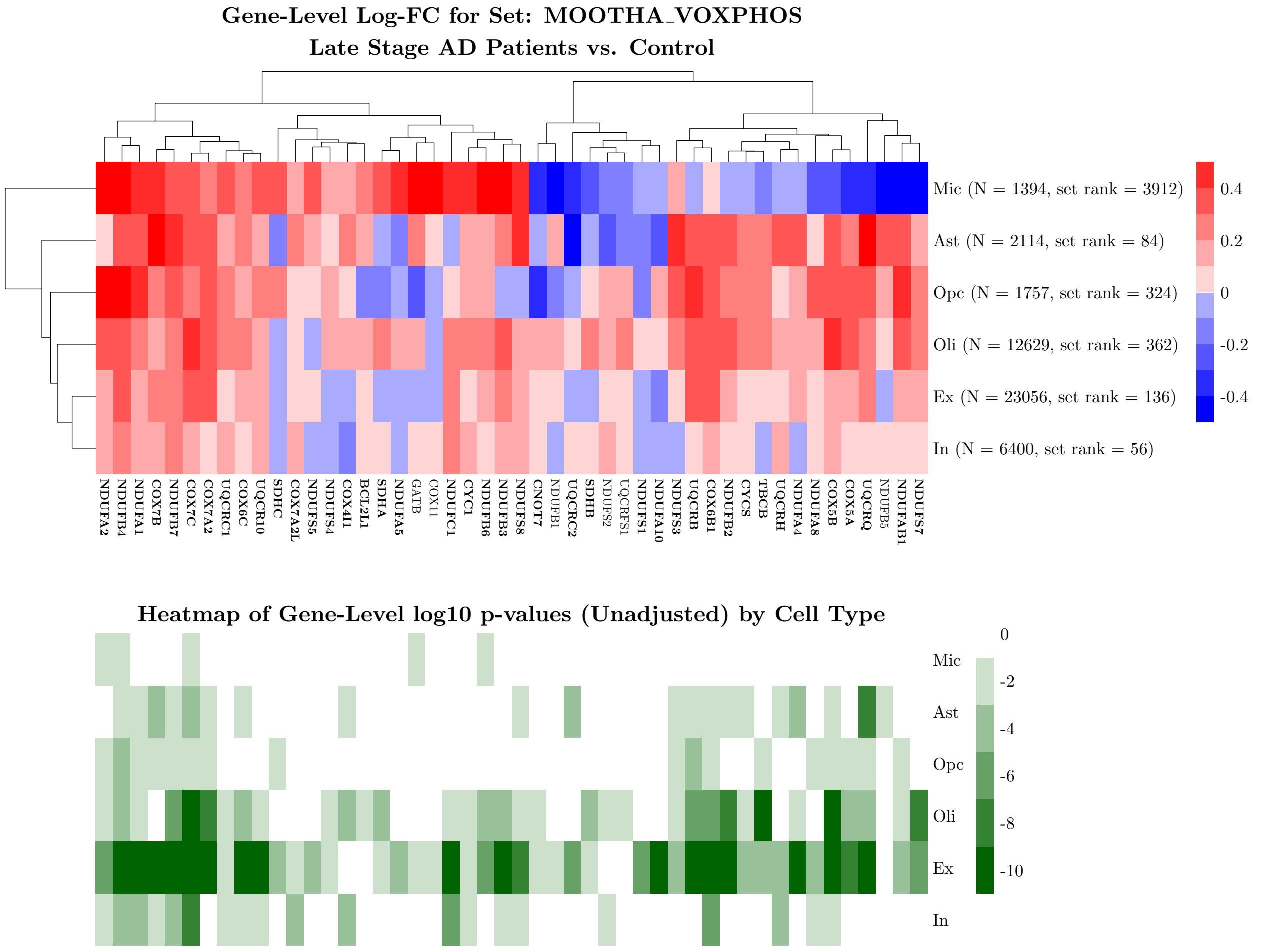


Supplementary Figure S20: **Variation in gene-level significance at the cell type level for genes in the MOOTHA_VOXPHOS pathway comparing late stage AD patients to controls**. Gene names that are bolded are signifcant over all cell types after FDR-adjustment of the Fisher's method $p$-value.

### S7 Additional Alzheimer's Results: Comparing AD Patients (Early and Late Stage) to Control

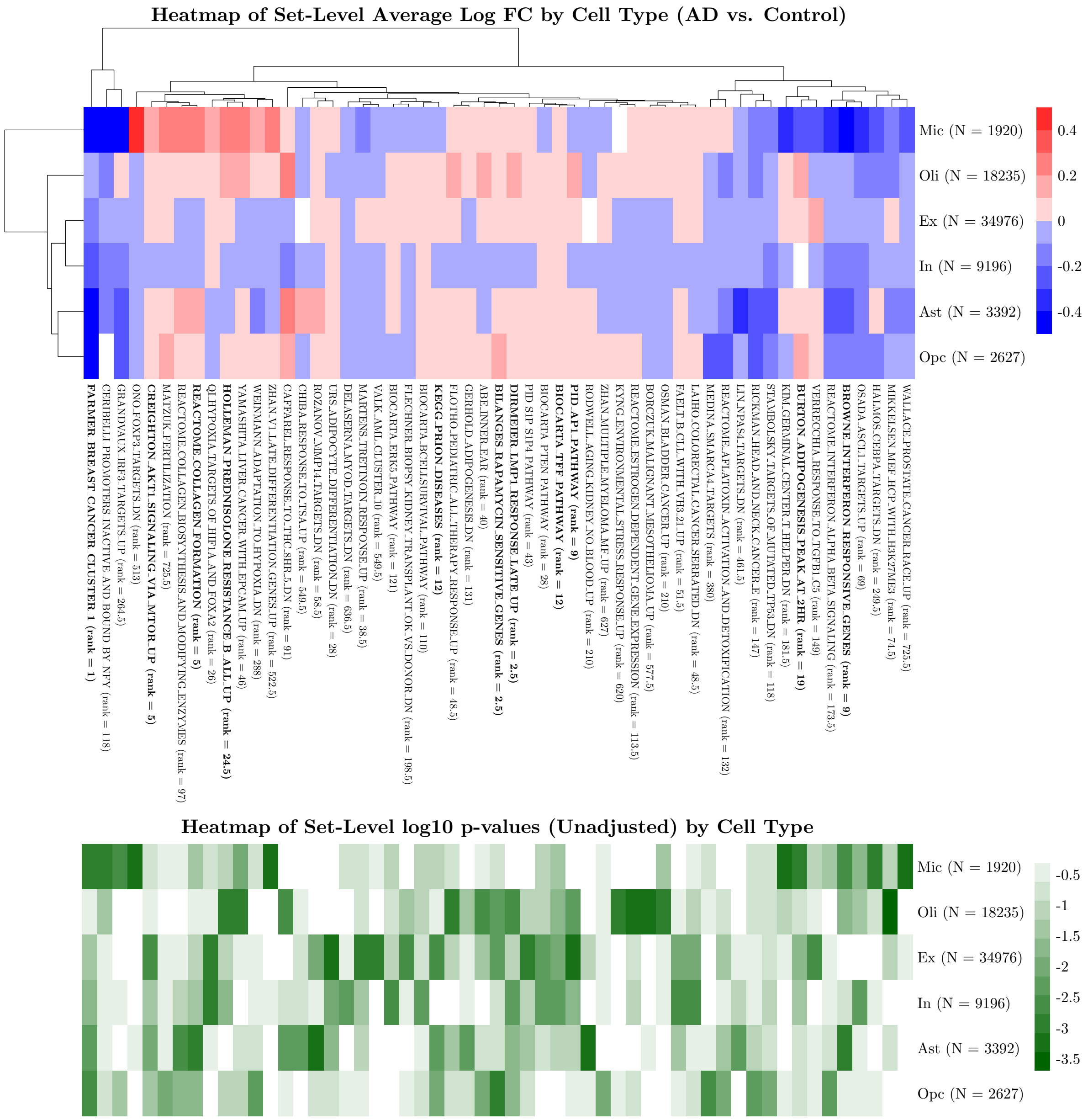


Supplementary Figure S21: **Heatmap of the most significant gene sets (and their corresponding p-values) comparing AD patients (early and late stage) to controls by cell type**. Sets plotted are among the top 10 in significance for at least once cell type. Sets in bold are significant over all cell types after FDR-adjustment of the Fisher's method $p$-value.

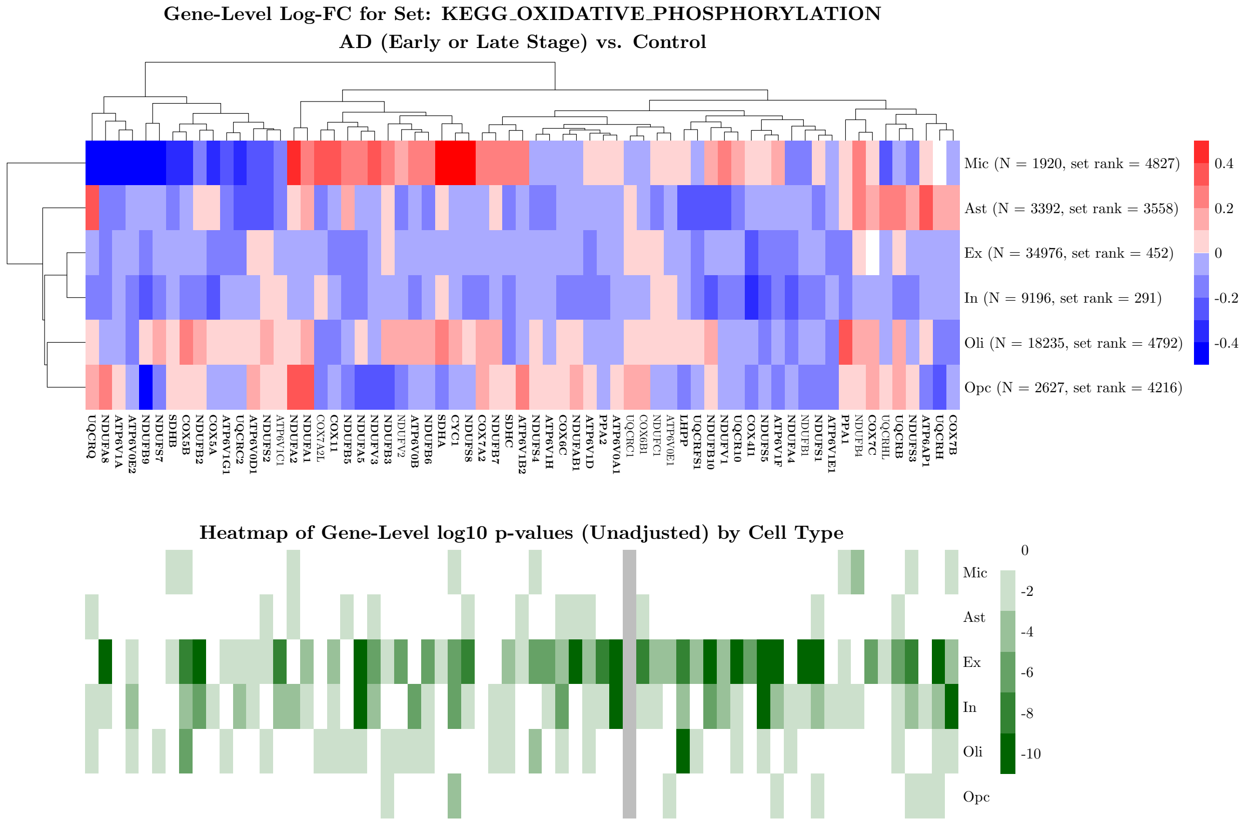


Supplementary Figure S22: **Variation in gene-level significance at the cell type level for genes in the KEGG_OXIDATIVE_PHOSPHORYLATION pathway comparing AD patients (early and late stage) to controls**. Gene names that are bolded are signifcant over all cell types after FDR-adjustment of the Fisher's method $p$-value.

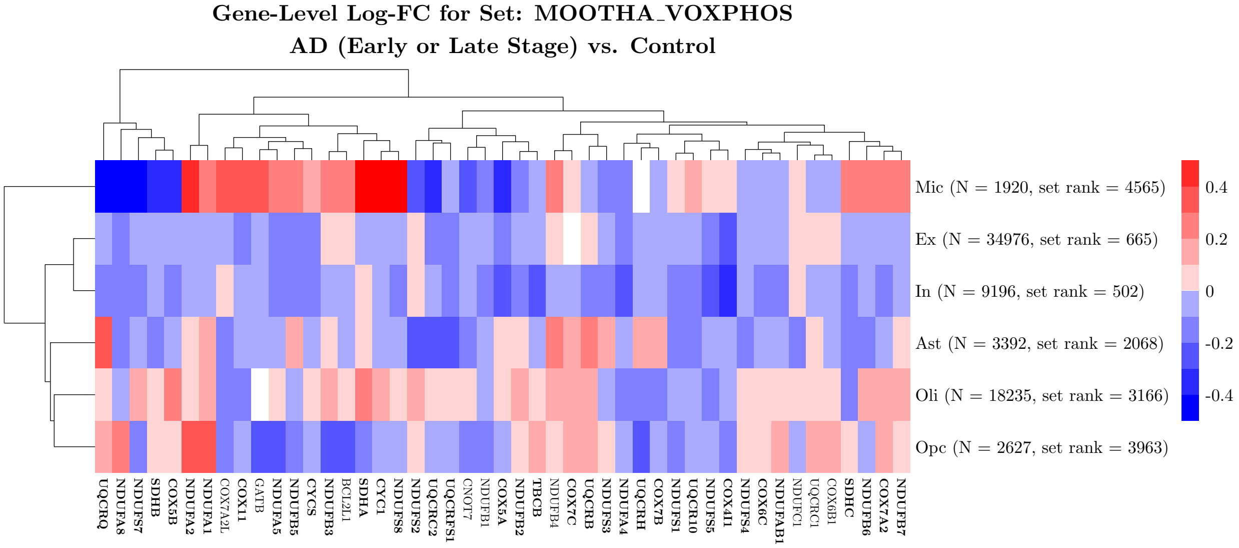


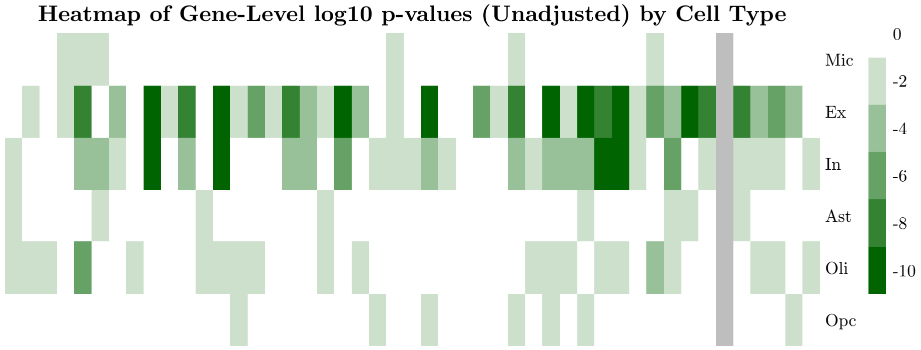


Supplementary Figure S23: **Variation in gene-level significance at the cell type level for genes in the MOOTHA_VOXPHOS pathway comparing AD patients (early and late stage) to controls**. Gene names that are bolded are signifcant over all cell types after FDR-adjustment of the Fisher's method $p$-value.

### S8 Additional Alzheimer's Results For Method Comparison

(A)

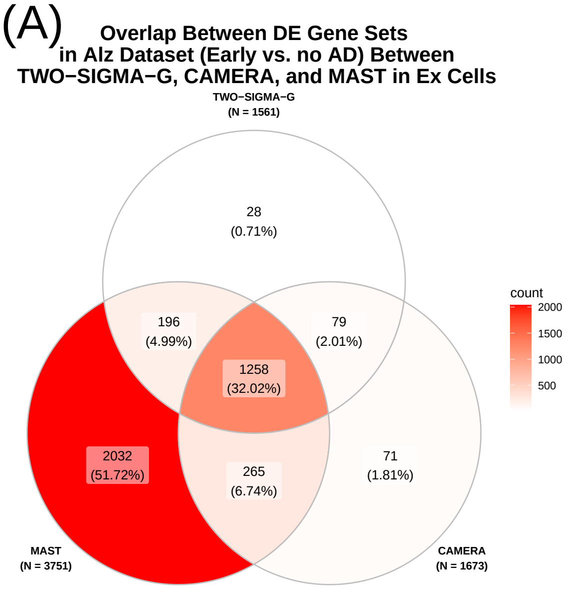


Supplementary Figure S24: **Venn diagram showing set-level overlap across methods among all DE gene sets before FDR adjustment for Ex cells in the Alzheimer's dataset (Early Stage AD vs. Control).**

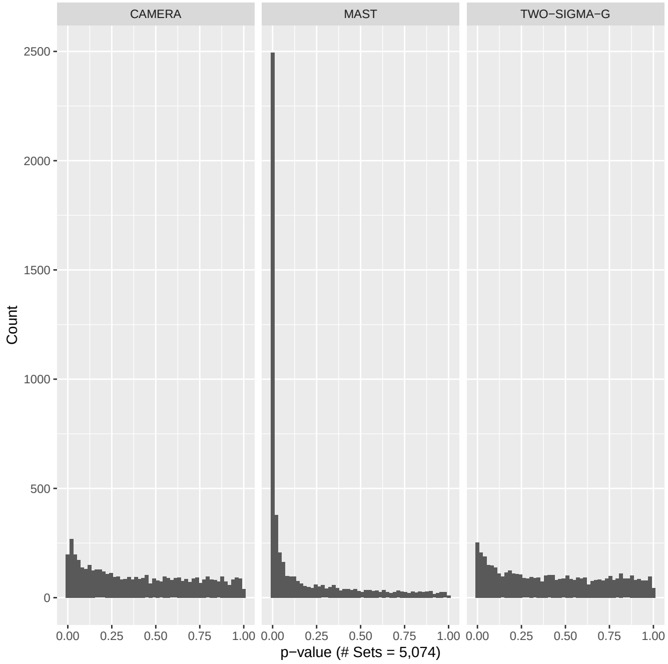


Supplementary Figure S25: **Histogram showing set-level $p$-values from Ex cells for MAST, TWO-SIGMA-G, and CAMERA in the Alzheimer's dataset (Early Stage AD vs. Control).**

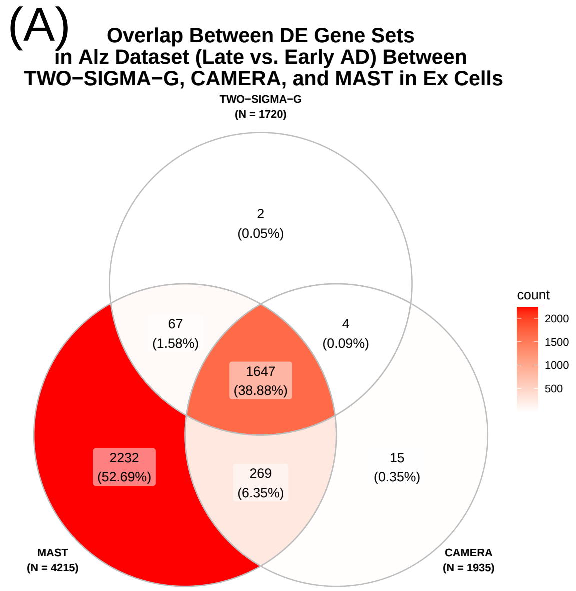


Supplementary Figure S26: **Venn diagram showing set-level overlap across methods among all DE gene sets before FDR adjustment for Ex cells in the Alzheimer's dataset (Late vs. Early Stage AD).**

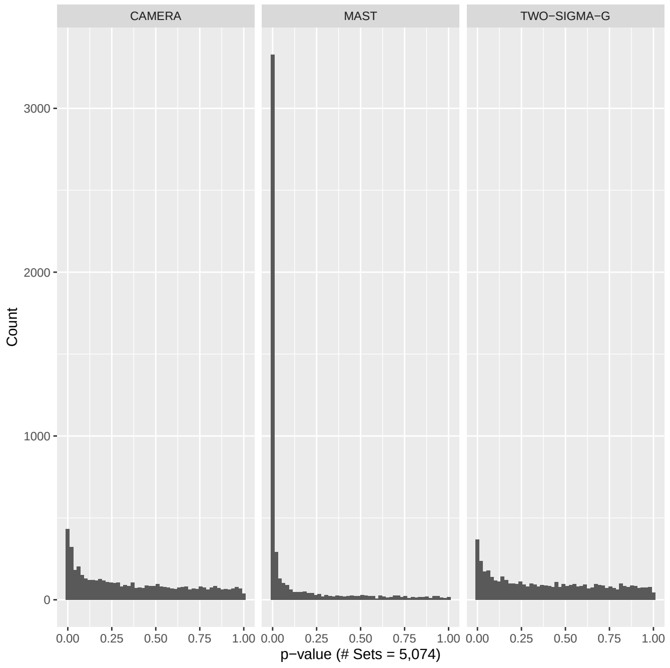


Supplementary Figure S27: **Histogram showing set-level $p$-values from Ex cells for MAST, TWO-SIGMA-G, and CAMERA in the Alzheimer's dataset (Late vs. Early Stage AD).**

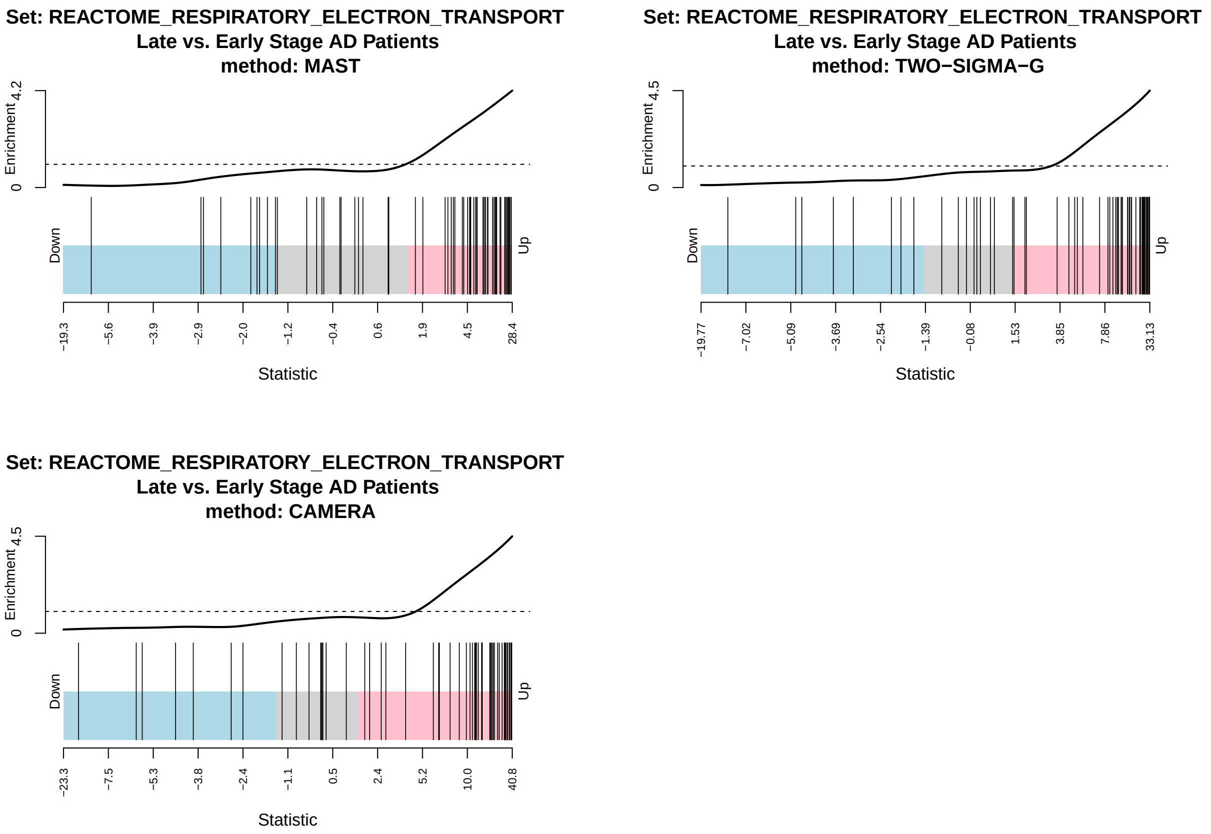


Supplementary Figure S28: **Barcode Plots for the Set REACTOME_RESPIRATORY_ELECTRON_TRANSPORT for MAST, TWO-SIGMA-G, and CAMERA comparing Late vs. Early stage AD.**

### S9 Random Effects in TWO-SIGMA-G in Both Datasets

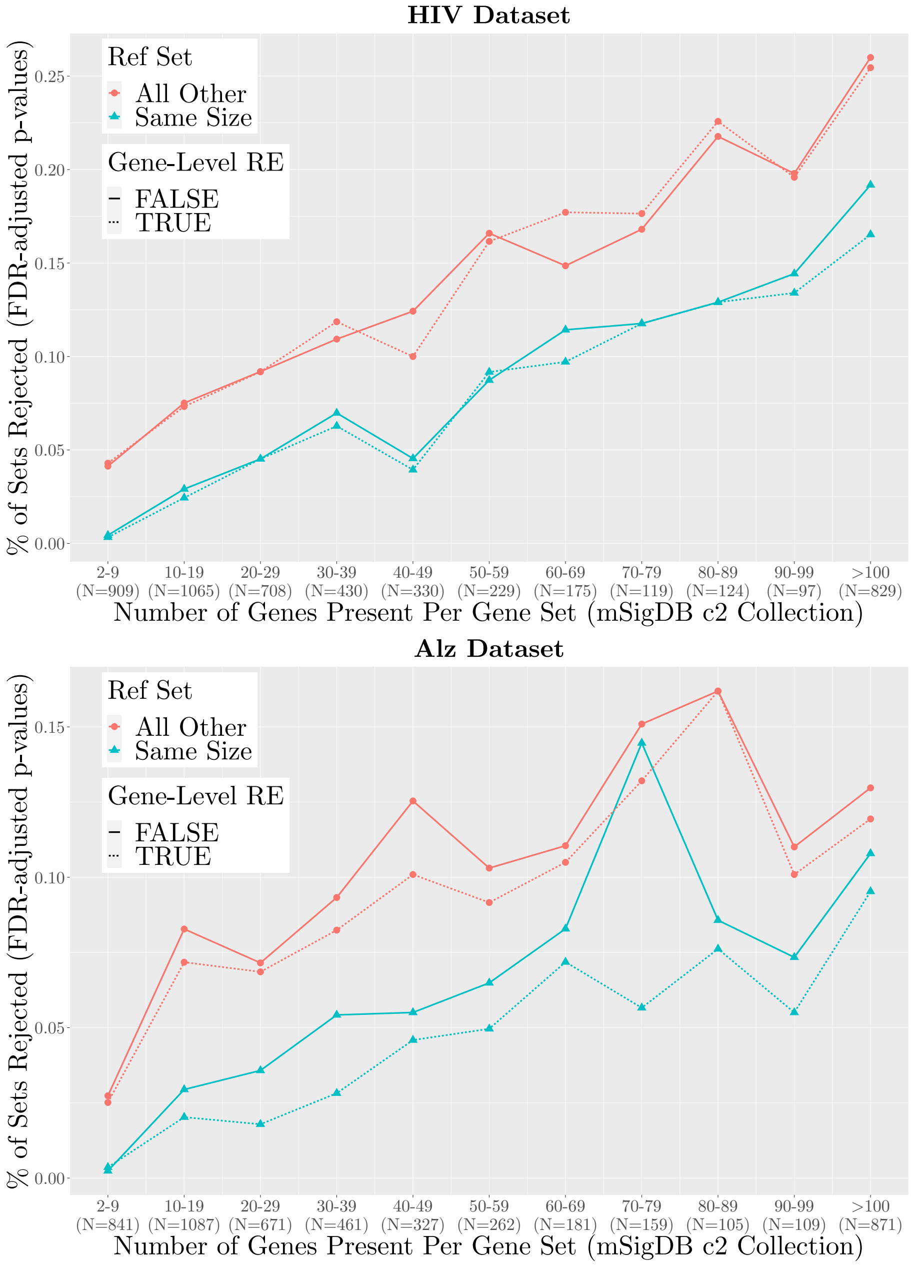


Supplementary Figure S29: **Percentage of sets rejected in the HIV and Alzheimer's datasets**. Fisher's-method p-values adjusted for FDR were used for testing in four settings varying the choice of reference set between the complement set of genes ("All Other") or a random reference of the same size as the test set ("Same Size"), and with and without random effects present at the gene-level. The presence of gene-level random effects in the model does not greatly affect the percentage of sets rejected in either the HIV dataset (top) or the Alzheimer's dataset (bottom).

#### S10 Relation between Gene-set Size and Expressed Genes in Both Datasets

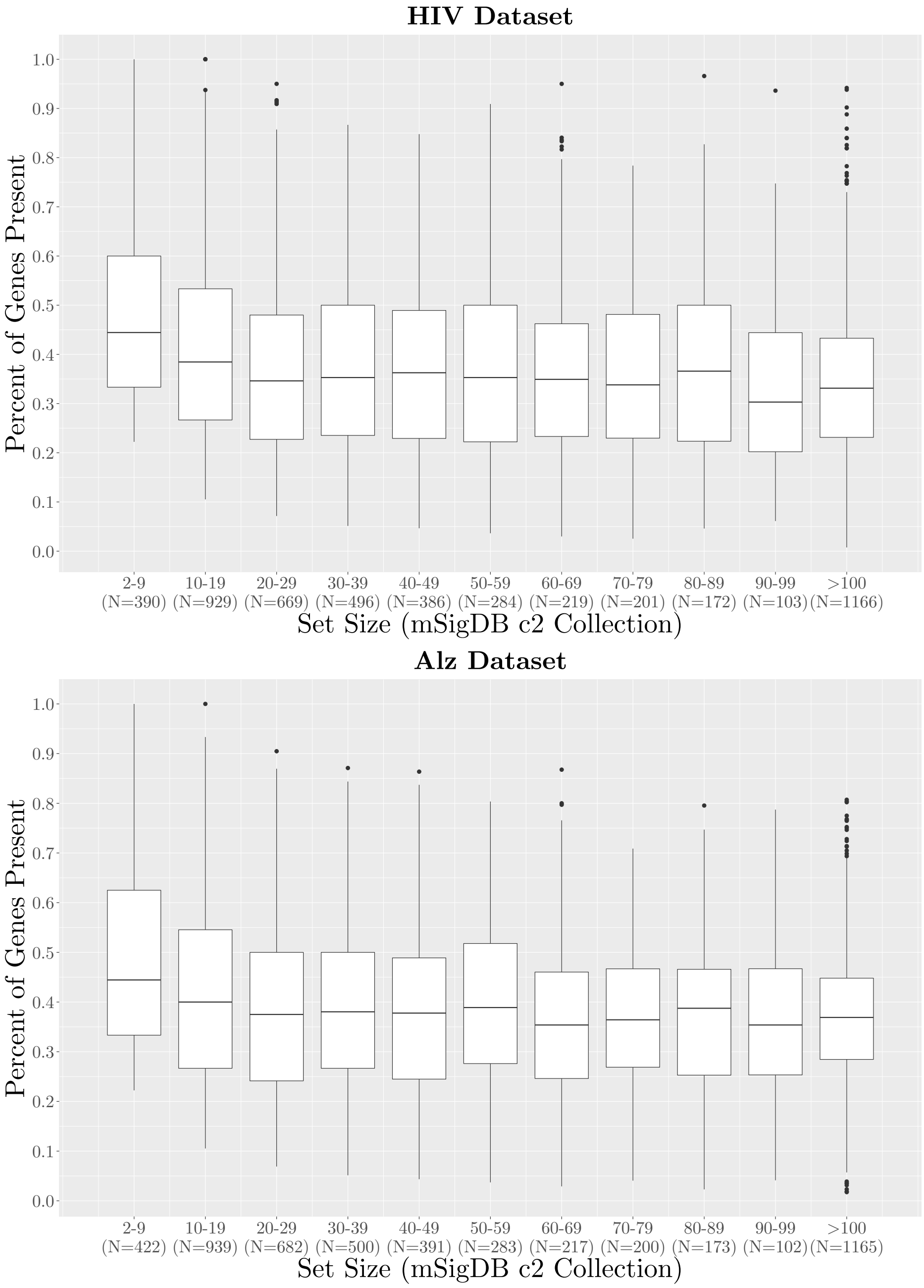


Supplementary Figure S30: **Percentage of genes present by set size in the HIV dataset (top) and the Alzheimer's dataset (bottom).**
